# Engineering Functionality Optimized fully human B7-H3 CAR T Cells for Enhanced Solid Tumor Therapy

**DOI:** 10.64898/2026.01.09.696281

**Authors:** Pradip Bajgain, Yang Feng, Mariela Puebla, Meijie Tian, Kuo-Sheng Hsu, Jaewon Lee, GuoJun Yu, Liping Yang, Steven Seaman, Mary Beth Hilton, Karen Morris, Niza Borchin, Jennifer D. Tran, Riley D Metcalfe, James C Cronk, Javed Khan, Anandani Nellan, Rosandra N Kaplan, Brad St. Croix

**Affiliations:** Tumor Angiogenesis Unit, Mouse Cancer Genetics Program, Center for Cancer Research (CCR), National Cancer Institute (NCI), National Institutes of Health (NIH), Frederick, MD 21702, USA; Pediatric Oncology Branch, CCR, NCI, NIH, Bethesda, MD, USA; Oncogenenomics Section, Genetics Branch, CCR, NCI, NIH, Bethesda, MD, USA; Basic Research Program, Leidos Biomedical Research Inc., Frederick National Laboratory for Cancer Research (FNLCR), Frederick, MD 21702, USA; Center for Structural Biology, CCR, NCI, NIH, Frederick, MD 21702, USA

**Keywords:** chimeric antigen receptor (CAR), adoptive T cell transfer, fully human antibody, phage display library, B7-H3, CD276, neuroblastoma, pancreatic ductal adenocarcinoma

## Abstract

B7-H3 is a cell surface protein overexpressed in many solid tumors and an attractive target for chimeric antigen receptor (CAR) T cell therapy. The most clinically advanced B7-H3 CARs are derived from murine monoclonal antibodies (mAbs) 376.96 and MGA271, which are now in phase **1/11** trials. However, non-human mAb sequences can provoke immune responses, leading to CAR T-cell rejection and therapeutic failure. Although scFv humanization reduces this risk, residual foreign residues within the variable domains remain. To overcome this limitation, we used in vitro phage display to generate fully human B7-H3-specific scFvs for CAR design. In pancreatic cancer, neuroblastoma, and glioblastoma xenograft models, CAR T cells incorporating the lead human binder Y111 were well tolerated and demonstrated superior antitumor activity compared with 376.96- and MGA271-based CARs. Y111 CAR treatment induced complete responses, tumor rejection, and significant survival benefits, identifying Y111 as a promising fully human B7-H3 CAR for solid tumors.

## INTRODUCTION

Chimeric antigen receptor (CAR) T cell therapy has achieved remarkable success in treatment of hematologic malignancies ^1^, prompting intensive efforts to extend this approach to solid tumors. B7-H3 (CD276) has emerged as a particularly promising target for CAR-based immunotherapy due to its high expression across diverse solid tumor types and limited expression in normal tissues^2^. B7-H3-specific CART cells using scFvs from 376.96 and MGA271 showed promise in preclinical studies^3^.4, leading to multiple phase **1/11** clinical trials worldwide^5^. Preliminary results suggest that targeting B7-H3+ tumors, including highly aggressive malignancies such as diffuse intrinsic pontine glioma (DIPG), is feasible, associated with manageable toxicities, and capable of achieving durable tumor regression in some patients^6^. However, as with other CAR T cell therapies directed against solid tumors, the expansion, persistence, and overall therapeutic efficacy of B7-H3-targeted CART cells remain suboptimal in most patients.

One underexplored factor that may substantially influence CAR T cell persistence and therapeutic efficacy is the immunogenicity of the single-chain variable fragment (scFv), which mediates antigen recognition7. Many CAR constructs incorporate scFvs derived from non-human monoclonal antibodies - most commonly murine - which can provoke immune-mediated clearance of the engineered T cells. Assessing how CAR immunogenicity affects clinical outcomes has been challenging due to variation in scFv sequences and heterogeneity of patient immune responses. Although these factors complicate the development of broadly applicable strategies to mitigate CAR immunogenicity, the use of fully human scFvs derived from phage display libraries offers a promising solution. Such libraries have proven to be reliable sources of therapeutic antibodies and have yielded scFvs used in antibody-based therapies and CARs targeting antigens such as CD19 and EGFRvlll^8^•^9^. Incorporating a fully human scFv into a CAR eliminates the need for humanization, reduces immunogenic potential, and may limit immune-mediated destruction of the infused T cells. Avoiding endogenous immune attack could enable improved CAR **T** cell proliferation and long-term persistence, thereby enhancing antitumor activity - particularly in solid tumors, which have historically shown resistance to CAR **T** cell therapies.

As with CARs targeting other tumor-associated antigens, most B7-H3-specific CARs developed to date, including the two currently in phase 1/11 U.S. trials, use murine-derived binders obtained through traditional hybridoma methods^10^ ^11^. Additional sources, such as murine scFvs and camelid single-domain antibodies isolated from phage display libraries, have also yielded B7-H3 binders suitable for CAR development^12,13^. However, because these binders originate from non-human species, they remain potentially immunogenic, posing a risk to CART cell persistence and therapeutic efficacy. Although scFv humanization can reduce immunogenicity, the approach is only partially effective, as foreign complementarity-determining regions (CDRs) and critical framework residues that cannot be fully humanized often retain immunogenic potential. To address these limitations, we generated a panel of B7-H3 CARs using scFvs isolated from a fully human antibody phage display library. This strategy eliminates the need for humanization and more effectively minimizes immunogenicity than approaches based on murine or camelid binders. In preclinical models of pancreatic cancer and pediatric neuroblastoma, one of these fully human CARs, Y111, exhibited robust and durable antitumor activity, outperforming murine- and camelid-derived CARs, including those currently under clinical investigation.

## RESULTS

### Fully Human scFv B7-H3 Binders Specifically Recognize Tumor-Expressed B7-H3

B7-H3 has emerged as an attractive target for CART cell-based cancer immunotherapy^3–5^. B7-H3 is overexpressed in multiple solid tumors including glioma, glioblastoma, pancreatic adenocarcinoma, and pediatric cancers such as rhabdomyosarcoma and neuroblastoma (Figure 1A), and elevated B7-H3 expression correlates with poor patient survival (Figure 1B). To engineer potent B7-H3-targeted CARs with minimal immunogenicity, we generated B7-H3-specific binders from a fully human scFv phage display library rather than using murine or other non-human antibodies. A na’fve human scFv library containing 8 x 10^10^ independent clones derived from B cells of 58 healthy donors was screened through three rounds of panning against biotinylated human B7-H3 (see Materials and Methods for details). While monoclonal phage ELISA identified hundreds of binders, DNA sequencing revealed that the majority derived from three repeatedly represented and distinct scFvs - Y868, Y111, and Y117 (Figure S1A). To assess specificity, we analyzed cell-surface binding of the three lead binders by flow cytometry. All three scFvs selectively recognized wild-type Panc1 pancreatic cancer cells but not B7-H3 knockout Panc1 cells (Panc1 B7-H3KO), confirming target-dependent recognition (Figures 1C-1D). Strong B7-H3 binding was also observed in naturally expressing HEK293T cells (Figure S1B) and in the pediatric neuroblastoma line IMRS (Figure 1E). Antigen sensitivity was evaluated by titrating scFv concentrations against a fixed number of IMRS cells. While all binders saturated at 10 µg/ml, Y111 exhibited markedly stronger binding at lower concentrations (Figure 1F), a pattern also observed in HEK293T cells (Figure S1C), indicating enhanced antigen sensitivity relative to Y868 and Y117. Collectively, these results demonstrate that all three fully human scFvs specifically recognize tumor-expressed B7-H3 without cross-reactivity to B7-H3-deficient cells, with Y111 displaying the highest antigen sensitivity.

**Figure 1.**
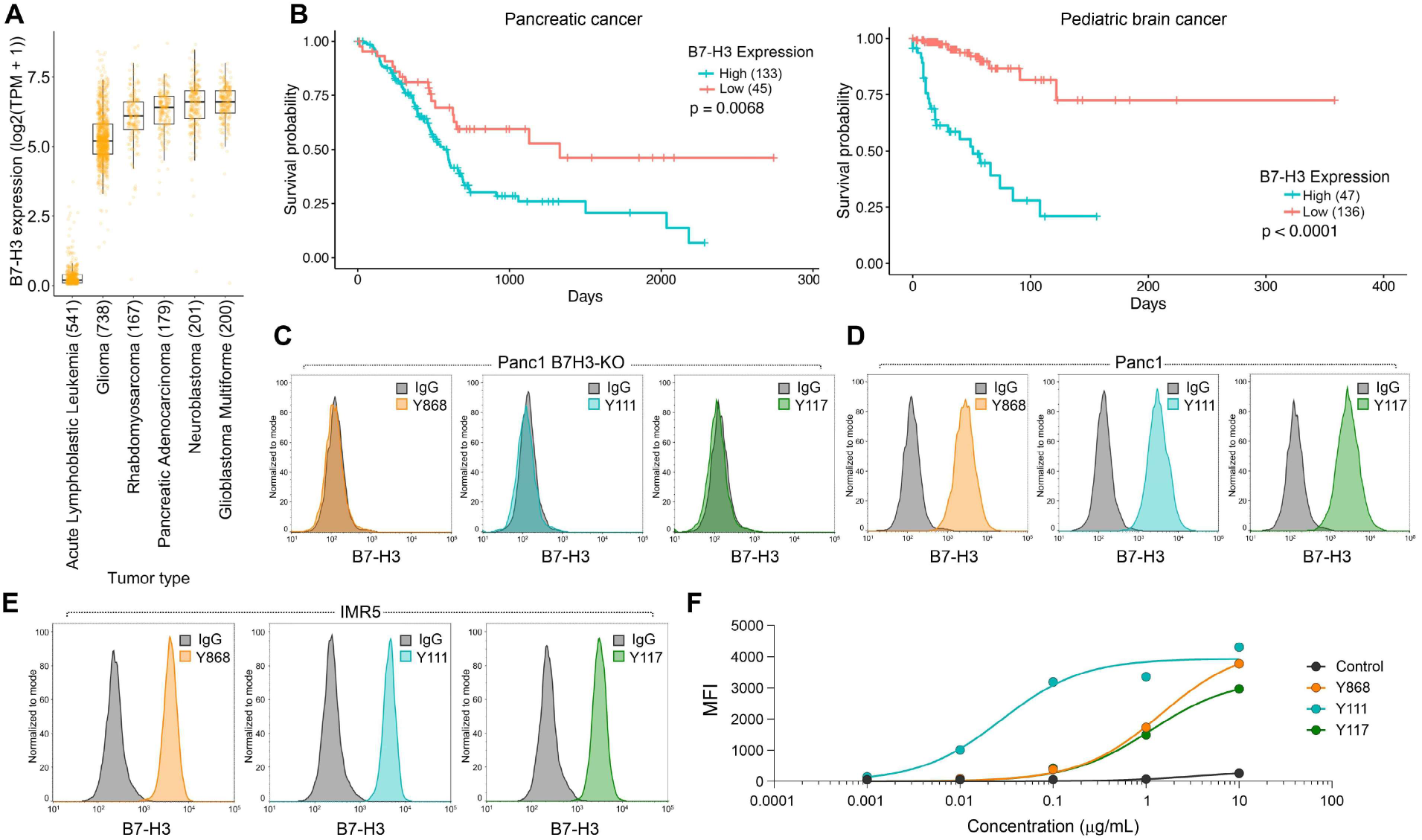
**B7-H3 overexpression in aggressive cancer types correlates with poor survival.** A) B7-H3 mRNA expression in aggressive human solid tumors from the DepMap Clinical Dataset. Acute Lymphoblastic Leukemia was included as a negative control. (B) Kaplan-Meier survival curves and log rank test for overall survival in B7-H3 high versus low expressing pancreatic and pediatric brain cancer from the TCGA database. n= 133 (B7-H3 high) and 45 (B7-H3 low) for pancreatic cancer, and n=47 (B7-H3 high) and 136 (B7-H3 low) for pediatric brain. P values were from a log-rank (Mantel-Cox) test. p = 0.0068 (Pancreatic cancer) and p < 0.0001 (Pediatric brain), high vs. low B7-H3. (C-E) Flow cytometry evaluating binding of Y868, Y111 and Y117 to Panc1 B7-H3KO (C), parent B7-H3+ Panc1 (B7-H3 positive) (D) or B7-H3+ IMR5 (E) cells. (F) Flow cytometry showing the dose-dependent binding of each of the scFvs to IMR5 cells. MFI: mean fluorescence Intensity.

### General Characteristics of T cells Engineered with the Fully Human B7-H3 CARs

Following validation of antigen specificity (Figures 1C-1F), second-generation CAR constructs were generated using each of the three human scFvs (Y868, Y111, Y117). These scFvs were fused to a previously optimized backbone comprised of CD28-derived hinge and transmembrane domains (28HTM), a 4-1BB co-stimulatory domain, and a CD3s signaling tail^14^ (Figure 2A). The CAR constructs, delivered via retroviral vectors, were transduced into activated primary human T cells, and surface CAR expression was assessed by flow cytometry three days post-transduction using an anti-human Fab antibody. All three CARs were efficiently expressed in the majority of T cells, whereas untransduced (UTD) T cells showed no detectable surface CAR expression (Figure 2B). As an alternative method to evaluate CAR surface expression and functionality, T cells were incubated with biotinylated recombinant human B7-H3 protein followed by APC-conjugated streptavidin. This antigen-binding assay produced results comparable to direct antibody staining for Y868 and Y111, and improved detection for Y117 (Figure 2C). To assess antigen sensitivity, recombinant B7-H3 protein was titrated against a fixed number of CART cells. Y111 CART cells demonstrated higher sensitivity, as indicated by increased frequencies of CAR+ cells and higher mean fluorescence intensity (MFI) at lower antigen concentrations relative to Y868 and Y117 (Figures 2D-2E), consistent with earlier scFv titration data (Figures 1F and S1C).

**Figure 2.**
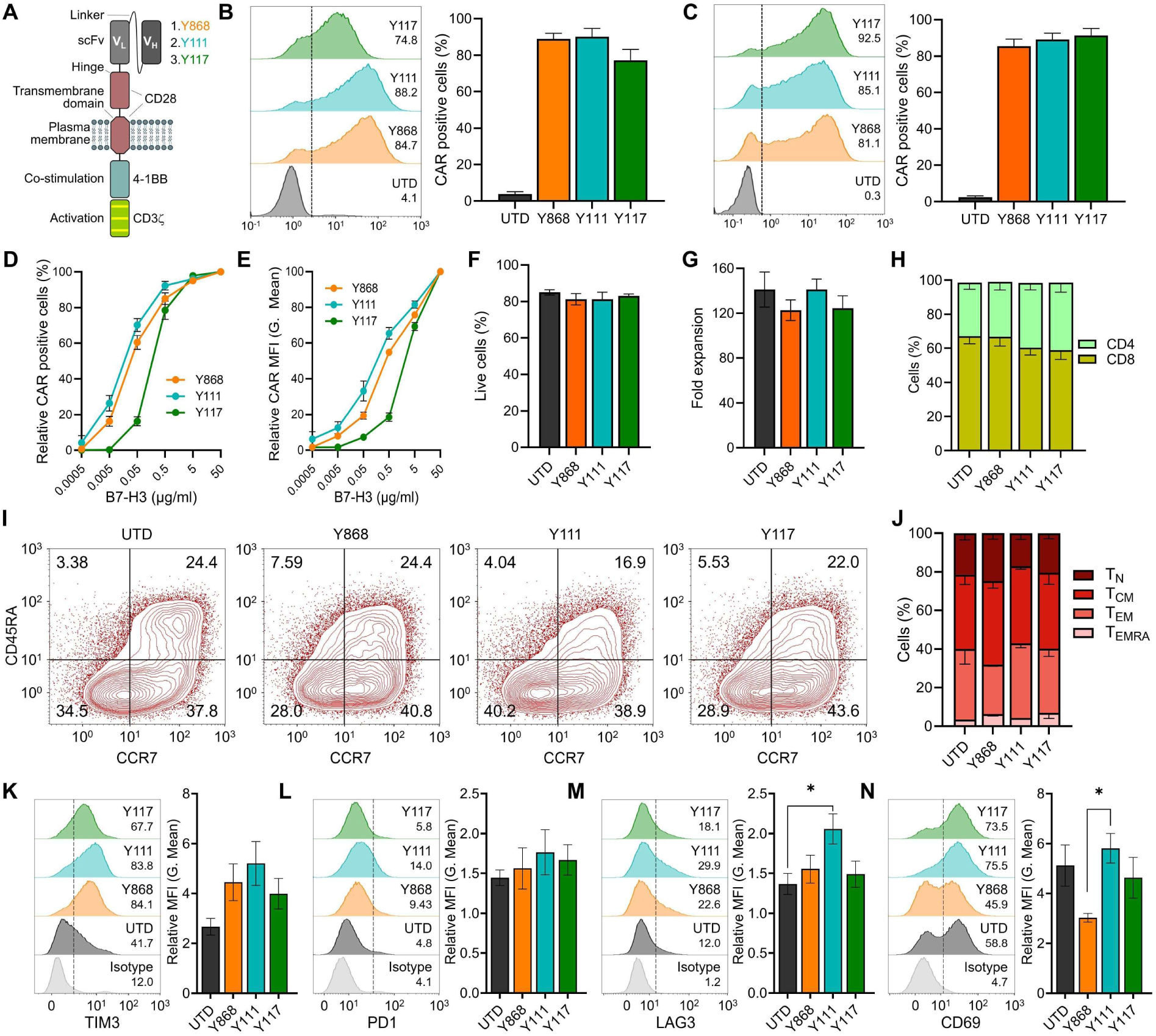
**Characterization of the T cell phenotype following B7-H3 CAR transduction.** (A) Schematic representation of CART structure used for each of the B7-H3 binders in this study. (B) Flow cytometry evaluation of CAR expression 72h post transduction using anti-human Fabs for detection. Left panel: representative histograms; Right panel: quantification of CAR expression. n=4 per group. (C) Flow cytometry evaluation of CAR expression 72h post transduction using recombinant biotinylated B7-H3 protein for detection. Left panel: representative histogram; Right panel: quantification of CAR expression. n=4 per group. (D) Flow cytometry showing biotinylated B7-H3 dose-dependent changes in % CAR positive T cells 72h post transduction. n=4 per group. (E) Flow cytometry showing biotinylated B7-H3 dose-dependent changes in MFI (geometric mean) of CAR positive cells 72h post transduction. n=4 per group. (F) Flow cytometry evaluation of T cell viability 11 days post CAR transduction. n=4 per group. (G) Fold T cell expansion 11 days post CAR transduction. n=4 group. (H) Flow cytometry analysis of CD4 and CD8 T cell fractions 10 days post CAR transduction. n=4 group. (I) Representative flow cytometry evaluation of T cell memory phenotype 10 days post CAR transduction. (J) Quantification of memory subset distribution from 4 independent donor samples including the example shown in (I). **(K-N)** Flow cytometry analysis of TIM3, PD1, LAG3 and CD69 levels in T cells 10 days post CAR transduction. Left panel: representative histogram; Right panel: quantification of MFI for each marker. n=4 per group.

We next examined the impact of CAR expression on T cell phenotype. No significant differences in cell viability or expansion were observed at day 11 post-transduction (Figures 2F-2G). The CD4:CD8 ratio was comparable to that of UTD cells, although all conditions showed a modest enrichment in cos+ T cells (Figure 2H). Memory subset distribution was also preserved, with central memory (TcM) and effector memory (TEM) cells predominating across all CAR and UTD groups (Figures 2I-2J). To evaluate T cell activation and exhaustion, we measured baseline expression of PD-1, TIM-3, LAG-3, and CD69 at day 11 post-transduction. Overall, CAR T cells displayed similar expression profiles to UTD cells, although Y111 CART cells exhibited elevated PD-1 levels (Figure 2L). Additionally, Y868 CAR T cells showed reduced CD69 expression compared with Y111 CAR T cells (Figure 2N). No other significant differences were detected among the groups (Figures 2K-2N). In summary, all three fully human CARs (Y868, Y111, Y117) were efficiently expressed in primary human T cells without affecting viability, subset distribution, or memory phenotype, with only modest changes in exhaustion and activation markers.

### B7-H3 CARs Selectively Kill B7-H3+Tumor Cells *In vitro*

To assess tumor specificity and cytotoxic activity of the fully human B7-H3 CAR T cells, we performed a luciferase-based killing assay. CAR-transduced or untransduced (UTD) T cells were co-cultured with GFP-firefly luciferase (GL)-tagged cancer cells at varying effector-to-target (E:T) ratios (5:1 to 0.25:1), and cytotoxicity was measured by luminescence after overnight incubation. As shown in Figure 3A, none of the CAR T cells exhibited cytotoxicity against B7-H3-negative Nalm6 leukemia cells or B7-H3-knockout NBEB neuroblastoma cells (NBEB B7-H3KO), confirming target-specific activity. In contrast, all three CART cell variants efficiently killed B7-H3+ pancreatic cancer (Panc1, HPAC, MIA PaCa-2) and neuroblastoma (NBEB, IMR5, LAN5) cells in an E:T ratio-dependent manner. Overall, each of the fully human B7-H3 CARs demonstrated comparable and potent on-target cytotoxicity against B7-H3+ tumor cells while sparing B7-H3-negative targets. (Figure 3A).

**Figure 3.**
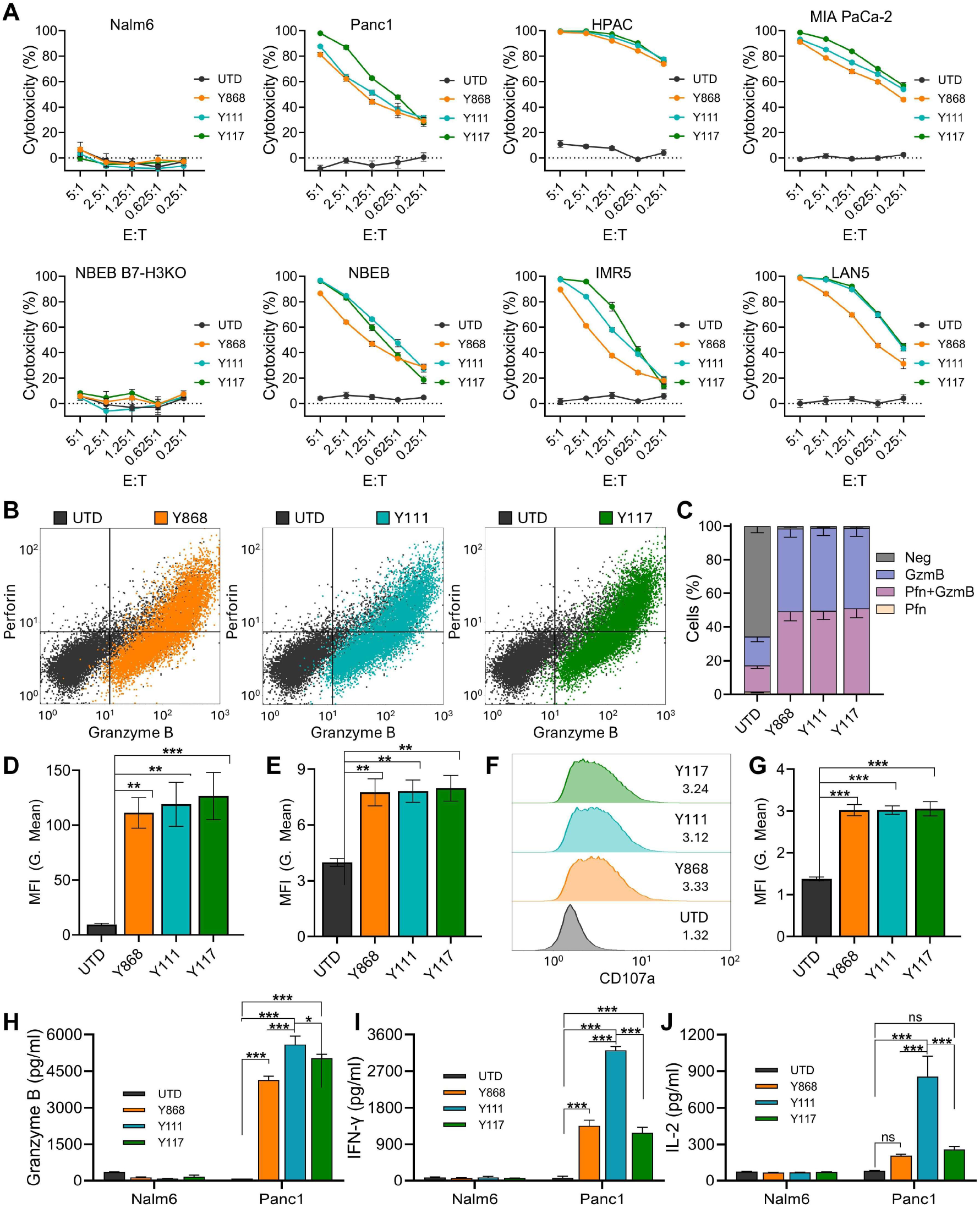
**B7-H3 CART cells show potent on-target cytotoxicity *in vitro*.** (A) Cytotoxic killing of luciferase-labelled tumor targets (T) by effector T cells (E) was monitored by co-culturing cells at various E:T cell ratios and measuring luminescence 48h later. B7-H3-Nalm6 and NBEB B7-H3KO cells were included as target negative controls. n=3 per group. (B) Representative flow cytometry evaluation of Perforin and Granzyme B expression in T cells 48h post co-culture with Panc1 tumor cells at E:T of 1:1. (C) Quantification of Perforin (Pfn) and Granzyme B (GzmB) % positive cells from four independent donor samples including the example shown in (B). (D) Quantification of Granzyme B MFI (geometric mean) from four independent samples including the example shown in (B). (E) Representative flow cytometry evaluation of CD107a in T cells 48h post co-culture with Panc1 tumor cells at E:T of 1:1. (G) Quantification of CD107a MFI from four independent samples including the example shown in (F). (H-J) ELISA measurement of Granzyme B (H), IFN-y (I) or IL-2 (J) production by T cells following co-culture with B7-H3- (Nalm6) or B7-H3+ (Panc1) cells for 48 hours. n=3 per group. Statistical significance was determined using a one-way ANOVA. *p:::.0.05, **p:::.0.01, ***p:::.0.001, ns, not significant.

To further evaluate effector function, we measured intracellular Granzyme Band Perforin levels following overnight co-culture with B7-H3+ Panc1 cells. All three CAR T cell populations showed robust induction of both cytolytic proteins, with >95% expressing Granzyme B alone or in combination with Perforin (Figures 3B-3C). The mean fluorescence intensity (MFI) of both proteins was significantly higher in CAR T cells compared with UTD controls, with no significant differences among the CAR constructs (Figures 3D-3E). Expression of CD107a (LAMP-1), a marker of degranulation, was similarly elevated in all CART cells compared to UTD cells, with comparable levels across the three CAR variants (Figures 3F-3G). Although these metrics indicated broadly similar cytolytic potential, Y111 CART cells elicited significantly higher levels of Granzyme B, interferon-gamma (IFN-y), and IL-2 in response to Panc1 cells compared with UTD and the other CAR groups (Figures 3H-3J). As expected, none of the T cells secreted these cytokines in response to B7-H3-negative Nalm6 cells, confirming antigen specificity. In summary, all three fully human B7-H3 CAR T cells mediated antigen-specific killing of B7-H3+ tumor cells and expressed key cytolytic proteins, while Y111 CAR T cells demonstrated enhanced effector cytokine secretion.

### B7-H3 CARs Regress Orthotopic Xenograft Pancreatic Tumors *In Vivo*

To evaluate the in vivo efficacy of fully human B7-H3 CAR T cells, we utilized an orthotopic xenograft model of pancreatic ductal adenocarcinoma (PDAC), a highly treatment-resistant cancer. 2.5×10^5^ luciferase-labeled Panc1 cells (Panc1-GL) were surgically implanted into the pancreas of NRG mice, followed by a single intravenous (i.v.) injection of CART cells two weeks later. Tumor progression was monitored weekly using bioluminescence imaging (BU), and body weight (BW) was tracked as a surrogate for treatment-related toxicity (Figure 4A). Mice receiving CAR T cells were carefully monitored for signs of toxicity, which was of particular concern for Y111 because of its cross-reactivity with murine B7-H3 (mB7-H3) (Figure S2A). No BW loss or other signs of toxicity were observed in any treatment group, indicating that the CAR T cell therapies were well tolerated (Figure S2B).

**Figure 4.**
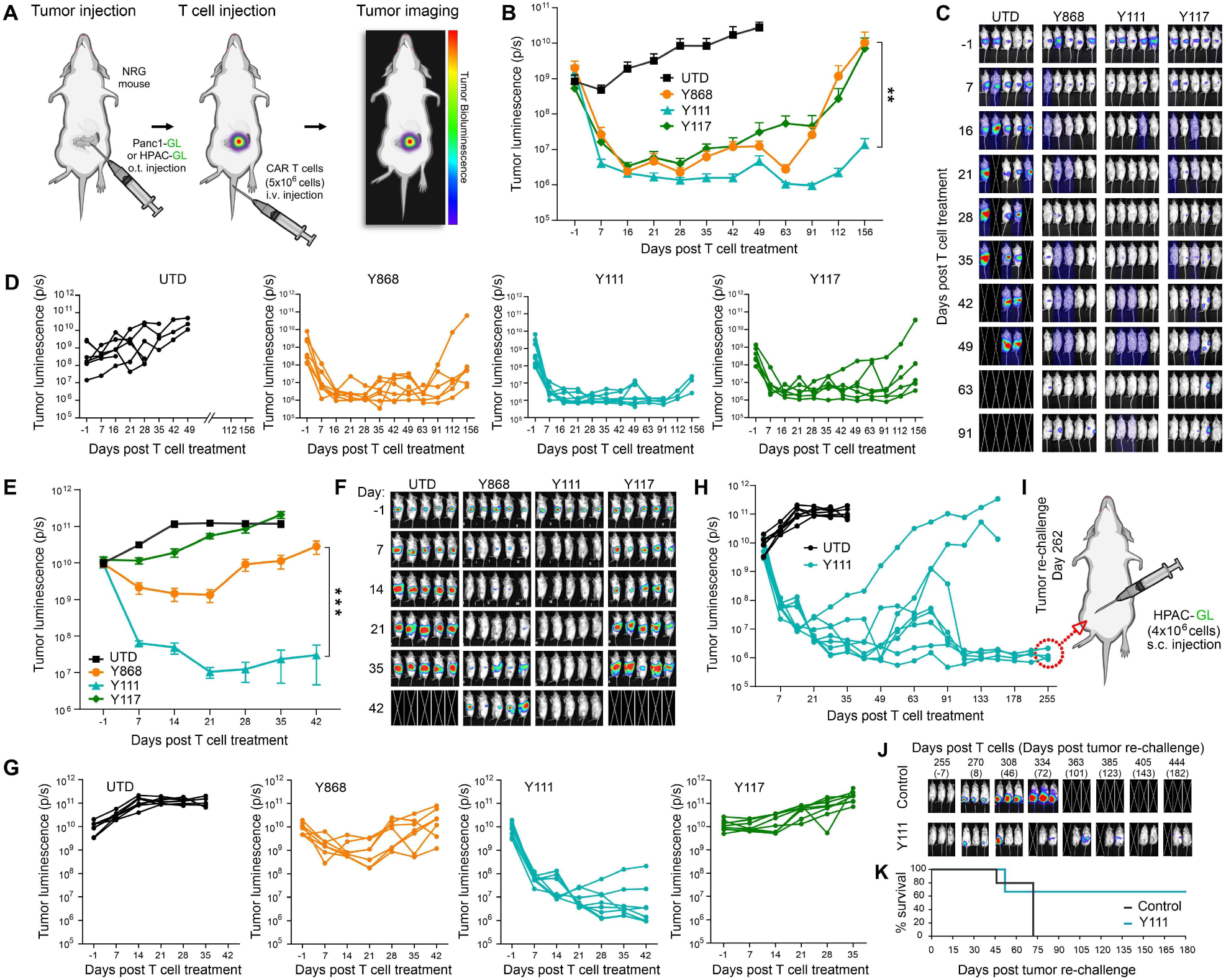
**Y111 B7-H3 CAR T cells display potent activity against orthotopic pancreatic tumors.** (A) Schematic overview of in vivo CAR T cell evaluation in orthotopic PDAC tumors. (B) BU was used to monitor growth of orthotopic Panc1-GL tumors. n=6-8 per group. (C) Images of representative mice from the study shown in (B). 5 mice shown per group. (D) Individual BU growth curves corresponding to the average growth curves shown in (B). (E) BU was used to monitor growth of orthotopic HPAC-GL tumors. n=7-8 per group. (F) Images of representative mice from the study shown in (E). 5 mice shown per group. (G) Individual BU growth curves corresponding to the average growth curves shown in (E). (H) Individual BU growth curves for the Y111 groups in E are shown up to day 255. The UTD control arm shown in G reached endpoint at day 35 and was also overlaid here for comparison. (I) Schematic overview of the s.c. rechallenge performed on long-term survivors in the Y111 group that reached day 255. (J) BU was used to monitor the growth of HPAC-GL tumors injected into surviving mice (n=3) in the Y111 group or age-matched naïve =5). 3 of the 5 control mice are shown. (K) Kaplan-Meier survival curves for the rechallenge study outlined in I and J. Statistical significance was determined using a log-rank (Mantel-Cox) test. n=5 (Control), 3 (Y111).

BU analysis revealed that all three CAR T cell variants induced substantial tumor regression within the first few weeks post-infusion (Figures 4B-4D). Over time, however, tumor relapses occurred earlier in mice treated with Y868 and Y117 CARs, whereas Y111 CAR T cells conferred more sustained tumor control (Figures 4B and 4D). To validate these findings, we tested the CARs in a second orthotopic PDAC model using luciferase-labeled HPAC-GL cells. lntrapancreatic HPAC tumors were established by surgical implantation of 2.0×10^5^ HPAC-GL cells, similar to the Panc1 model (Figure 4A). T cells were administered i.v. 12 days after the surgery. As in the Panc1 model, no BW loss was observed following CAR T cell infusion. Some BW reduction, particularly in the UTD group, was noted and attributed to progressive tumor burden (Figure S2C). In this aggressive model of PDAC, only Y111 CART cells achieved robust and durable tumor regression, whereas Y868 and Y117 delayed progression without providing lasting control (Figures 4E-4G). Between the two less effective CARs, Y868 displayed superior anti-tumor activity relative to Y117. All mice treated with UTD, Y868, or Y117 T cells reached humane endpoints by day 42 post-infusion due to tumor burden. In contrast, Y111 CAR T cells produced long-term survival in most mice: six of eight survived at least 100 days, with only two experiencing tumor recurrence requiring euthanasia (Figures 4H-4I). Despite two additional deaths, the cause of which was unclear, four of eight Y111-treated mice (50%) remained alive and tumor-free at day 255 (∼8.5 months) post-treatment, demonstrating durable therapeutic benefit (Figure 4H). In summary, all three B7-H3 CART cells showed initial antitumor activity in orthotopic PDAC models; however, Y111 CAR T cells provided superior, long-lasting tumor control, minimal toxicity, and the greatest survival benefit.

### Y111 CART Cells Confer Long-Term Immunity Against Tumor Rechallenge

To evaluate the persistence and long-term antitumor function of Y111 CART cells, we performed a tumor rechallenge in the surviving mice from the HPAC orthotopic model (Figure 4E). On day 262 post-infusion, HPAC-GL cells were injected subcutaneously (s.c.) at a site distant from the pancreas in three Y111-treated cured mice. Five age-matched, tumor- and T cell-na’ive NRG mice received identical s.c. tumor implants as controls (Figure 41). Remarkably, BU monitoring revealed delayed or absent tumor engraftment in two of the three rechallenged Y111 mice, whereas all five control mice exhibited rapid tumor growth (Figure 4J). One rechallenged mouse developed progressive disease and required euthanasia, similar to all control animals. The remaining two mice maintained low tumor burdens for over six months following rechallenge (Figure 4K). On day 444 (∼15 months post-CART cell treatment), both long-term survivors were euthanized due to age-related severe health deterioration, having successfully resisted both the primary orthotopic tumor and the secondary s.c. tumor. These findings demonstrate that Y111 CAR T cells mediate potent and durable PDAC regression and can persist long-term to provide protective immunity against tumor re-exposure. Among the three fully human B7-H3 CAR constructs tested, Y111 consistently displayed superior efficacy and sustained functional persistence.

### Comparison of Y111 with Advanced CARs Derived from Murine and Camelid Binders

Most reported B7-H3-targeting CARs have been generated using murine-derived anti-B7-H3 monoclonal antibodies (mAbs), including scFv-based constructs derived from 376.96 and MGA271 mAbs^10^•^11^, both currently under evaluation in multiple US clinical trials. More recently, camelid-derived single-domain antibodies have also been explored for CAR development^12^. To directly compare the efficacy of our fully human CARs - particularly Y111 - with these advanced non-human CARs, we generated three benchmark constructs using antibodies from 376.96, MGA271, and 812 (a camelid-derived antibody^12^), all fused to a common 28HTM-41BB-CD3Z: backbone (Figure 2A). The specificity and binding kinetics of the benchmark antibodies have been described previously^3^•^4^ and were independently validated in our laboratory (Figure S3). To compare CARs derived from human and non-human antibodies, we first evaluated Y868, Y117, Y111, and 376.96 CARs in a disseminated neuroblastoma model established via intravenous injection of IMR5-GL cells (Figure 5A). At a suboptimal dose of 2 million CAR T cells, Y868 and Y117 CARs failed to elicit therapeutic benefit, whereas Y111 CAR T cells induced rapid tumor regression, outperforming the 376.96 CAR (Figures 58-5D). These responses were transient, reflecting the deliberately challenging low-dose conditions used to differentiate CAR performance. Given the consistent superiority of Y111 over Y868 and Y117, we next compared Y111 to 376.96 and MGA271 CARs in a sub-optimally dosed rhabdomyosarcoma model. Mice bearing intramuscular JR-1 tumors received 2 million UTD, Y111, 376.96, or MGA271 CART cells (Figure S4). Y111 CART cells again demonstrated the strongest anti-tumor activity, followed by 376.96 CAR, whereas MGA271 showed minimal or no efficacy (Figures S4A-S4E), translating into significantly improved survival for Y111 CAR T-treated mice (Figure S4F).

**Figure 5.**
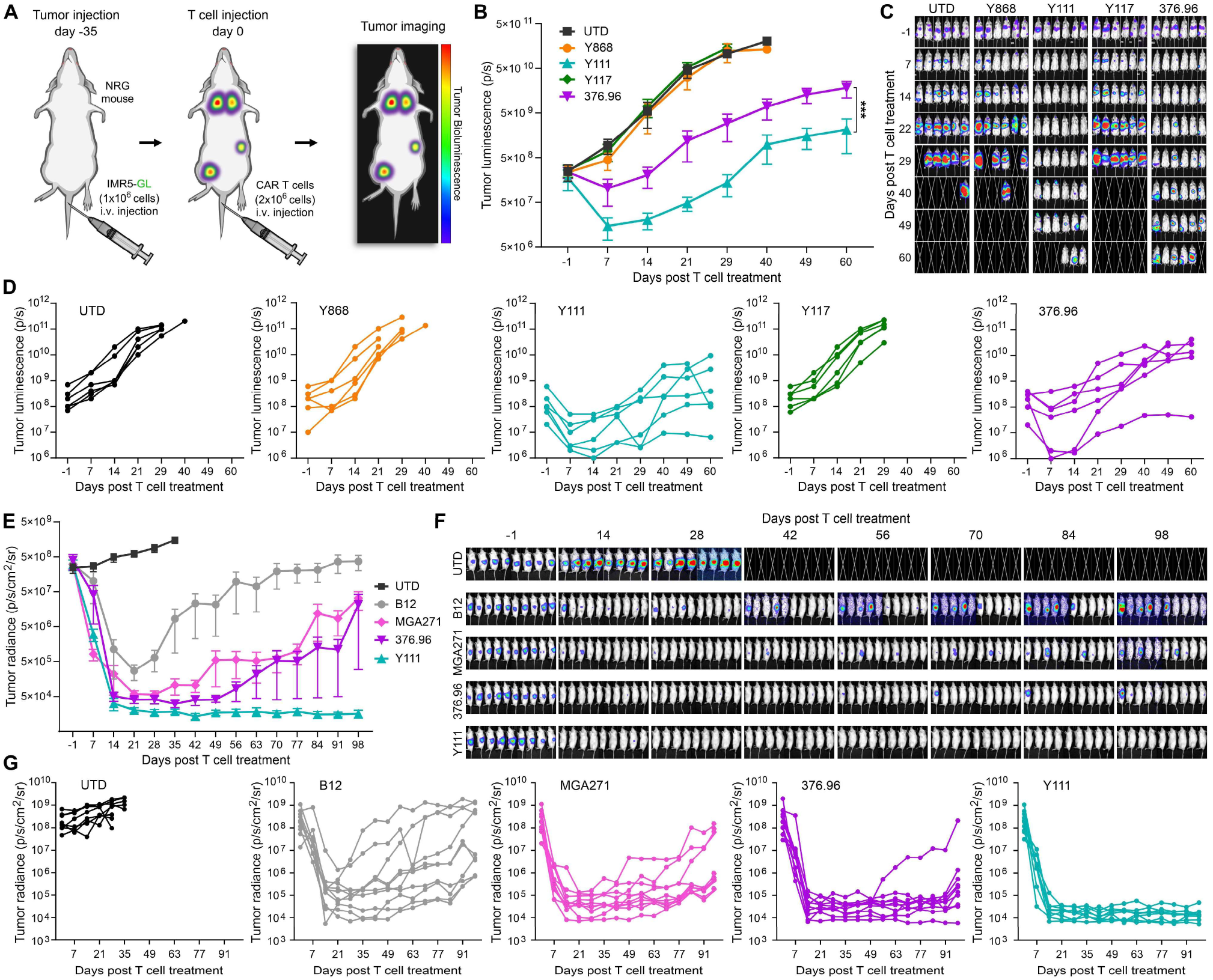
**Y111 compares favorably to B7-H3 benchmark CARs.** (A) Schematic overview of the disseminated neuroblastoma CART study. (B) BU was used to monitor growth of orthotopic IMR5-GL tumors. n=6 per group. (C) Images from the study shown in (B). (D) Individual BU growth curves corresponding to the average growth curves shown in (B). (E) BU was used to monitor growth of orthotopic Panc1-GL tumors. n=8-10 per group. (F) Images from the study shown in (E). (G) Individual BU growth curves corresponding to the average growth curves shown in (E).

Next, we compared four CARs - Y111, 376.96, MGA271, and B12 - in the orthotopic (o.t.) Panc1 PDAC model (described in Figure 4). All CAR T cell treatments initially reduced tumor burden (Figures 5E-5G). However, responses in the B12 group were incomplete and heterogeneous, and tumors rapidly recurred in MGA271-treated mice despite early regression. The 376.96 CAR outperformed MGA271 and B12 but still resulted in late relapses in most mice. In contrast, Y111 CART cells achieved the most consistent, complete, and durable tumor control across the cohort (Figures 5F-5G). In summary, the Y111 CAR consistently outperformed CARs constructed from murine and camelid-derived binders across multiple tumor models, demonstrating superior and durable anti-tumor efficacy and positioning it as a strong candidate for clinical development.

### Y111 CAR Demonstrates Superior Efficacy Against Bulky Neuroblastoma Tumors

Subcutaneous tumors represent a stringent therapeutic challenge due to their large mass, poor vascularization, and enrichment of T cell-inhibitory factors. To evaluate the anti-tumor activity of Y111 CAR T cells under these conditions, IMR5 neuroblastoma cells were implanted subcutaneously in NRG mice, followed by intravenous administration of CART cells (Figure 6A). In this aggressive model, B12 CAR T cells induced only minimal delays in tumor progression compared to the UTD (Figures 6B-6C). MGA271 CAR T cells slowed tumor growth but were unable to induce regression (Figure 6D). Responses in the 376.96 CAR group were heterogeneous: roughly half of the mice showed no tumor control, whereas the remainder exhibited transient stabilization lasting 3-4 weeks post-infusion (Figure 6E). In contrast, Y111 CAR T cells elicited the most robust anti-tumor responses, consistent with their superior performance in other tumor models (Figure 5). Although 3 of 10 mice did not respond to treatment, the remaining animals demonstrated marked tumor regression (Figure 6F). Notably, some tumors larger than 1000 mm^3^ regressed following Y111 CAR infusion, resulting in significantly prolonged survival compared with other groups (Figure 6G).

**Figure 6.**
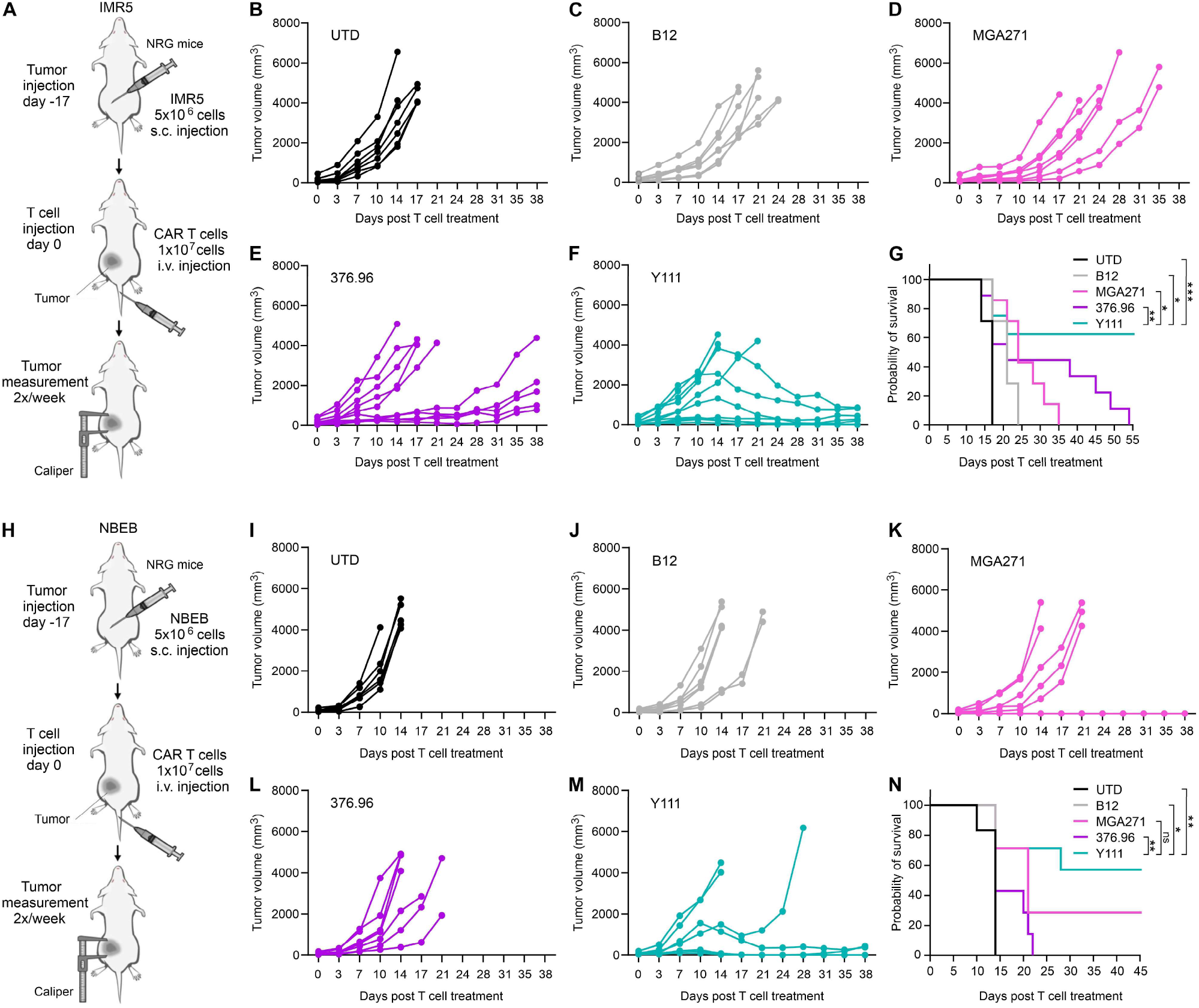
**Y111 shows potent activity against aggressive subcutaneous solid tumors.** (A) Schematic overview of the CAR T study targeting s.c. IMR5 neuroblastoma. (B-F) Individual tumor growth curves were monitored using calipers following i.v. injection of UTD (B), B12 (C), MGA271 (D), 376.96 (E) or Y111 (F) CART cells. n=7-10 per group. (G) Kaplan-Meier survival curves for the study outlined in A to F. Statistical significance was determined using a log-rank (Mantel-Cox) test. n=7 (UTD, B12, and MGA271) and 10 (Y111 and 376.96) (H) Schematic overview of the CART study targeting s.c. NBEB neuroblastoma. (I-M) Individual tumor growth curves were monitored using calipers following i.v. injection of UTD (I), B12 (J), MGA271 (K), 376.96 (L) or Y111 (M). n=6-7 per group. (H) Kaplan-Meier survival curves for the study outlined in H and I. Statistical significance was determined using a log-rank (Mantel-Cox) test. n=6 (UTD) and 7 (B12, MGA271, 376.96, and Y111)

We further validated these findings in a second subcutaneous neuroblastoma model using NBEB cells (Figure 6H). Consistent with the IMR5 results, B12, MGA271, and 376.96 CART cells conferred minimal to no therapeutic benefit. Nearly all mice in these groups required euthanasia by day 21, with only a single MGA271-treated mouse exhibiting measurable tumor regression (Figures 6I-6L). In contrast, the majority of mice treated with Y111 CAR T cells (4 of 7) demonstrated clear tumor regression accompanied by improved survival (Figures 6M-6N). Collectively, these results corroborate our prior observations and demonstrate that Y111 CART cells consistently outperform MGA271-, 376.96-, and B12-based CARs in aggressive subcutaneous models of pediatric neuroblastoma.

B7-H3 is an emerging target antigen in brain tumors, and B7-H3-directed CAR T cells have shown early signs of clinical activity and safety in both pediatric and adult patients^15–18^. Glioblastoma highly express B7H3 and higher expression is associated with poor prognosis in pediatric brain tumors (Figures 1A-1B). To evaluate the efficacy of Y111 CAR against glioblastoma, we established orthotopic GBM01 tumors in NRG mice and administered UTD, Y111, 376.96, MGA271, or B12 CAR T cells intravenously (Figure S5A). Consistent with our findings in other tumor models, Y111 CAR T cells achieved superior tumor control in this aggressive GBM setting, whereas other CARs exhibited only marginal or no activity (Figures S5A-S5F). This enhanced anti-tumor activity translated into a significant survival benefit, which was not observed with the comparator CARs (Figure S5G).

The enhanced activity of the Y111 CAR raises an important question: why is it more potent in vivo than the benchmark CARs MGA271 and 376.96? Although the precise mechanism remains under investigation, several distinctive features of the Y111 antibody suggest plausible explanations. Because prior studies have shown that epitope location can profoundly influence CAR function^19^•^20^, we first compared the binding sites of MGA276, 376.96 and Y111 on B7-H3. Domain-swap experiments between B7-H3 and its closest homolog, PD-L1, revealed that Y111 uniquely requires both the V1 and C1 domains for optimal affinity, indicating recognition of a discontinuous epitope spanning these regions (Figures S6A-S6B). Y111 also demonstrated greater thermal stability than MGA271 and 376.96, resisting denaturation at temperatures below 70°C (Figure S6C). Finally, when recombinant human B7-H3 extracellular domains were incubated with increasing concentrations of antibody, Y111 predominantly formed 1:1 complexes, whereas MGA271 and 376.96 generated higher-order aggregates (Figure S6D-S6E). This observation is notable given that CARs can be activated not only by membrane-bound antigen but also, under certain conditions, by soluble antigen capable of cross-linking and dimerizing CAR molecules^21^. Because the human 4Ig B7-H3 ectodomain contains an internal repeat^22^•^23^ and can be shed into the circulation of cancer patients^24^•^2^6,many B7-H3 CARs may be susceptible to tonic signaling driven by bivalent or multivalent cross-linking in solution. By contrast, the unusual monomeric binding preference of Y111 (Figures S3A and S6D-S6E) may reduce susceptibility to such tonic activation, ensuring that CAR clustering - and thus signaling - occurs predominantly upon engagement with B7-H3 on target cells.

## DISCUSSION

B7-H3 is broadly overexpressed across a range of solid tumors, including several with limited therapeutic options, making it an attractive target for CART cell immunotherapy^2^•^5^. Yet, despite the transformative success of CAR T therapies in hematologic malignancies, their activity in solid tumors remains inconsistent. Multiple factors contribute to this challenge, and one increasingly recognized determinant of CAR performance is the antibody binder incorporated into the receptor. In this study, our goal was to identify a fully human B7-H3-specific scFv with superior functional properties compared to the murine-derived binders used in clinical-stage CARs. Leveraging a large phage display library of fully human scFvs, we isolated several unique B7-H3 binders and prioritized three, Y868, Y111, and Y117, for detailed evaluation based on their potent tumor-cell recognition and cytotoxic activity in vitro. To ensure that observed functional differences were attributable solely to the antibody binder, all CARs were constructed using an identical architecture consisting of a CD28-derived hinge/transmembrane domain, a 4-1BB costimulatory domain, and a CD3s activation domain. This optimized backbone has previously demonstrated strong performance in CARs targeting CD19, HER2, and B7-H3^1^4, enabling a controlled comparison across binders. Across multiple xenograft models, including pancreatic ductal adenocarcinoma, neuroblastoma, glioma, and rhabdomyosarcoma, all three fully human B7-H3 CART cells mediated substantial anti-tumor activity. Among them, Y111 emerged as the most potent, inducing deep and durable tumor regressions and demonstrating long-term functional persistence, as evidenced by resistance to tumor rechallenge more than 15 months after initial treatment. Importantly, the Y111 CAR also outperformed benchmark CARs derived from the murine 376.96 and MGA271 antibodies, both of which are currently undergoing clinical evaluation, highlighting the therapeutic advantage of this newly identified human binder.

Murine-derived antibodies remain the predominant source of CAR binders, including in the most clinically advanced B7-H3-targeted CARs. Camelid-derived single-domain antibodies are also being explored^12^. While these non-human sources can generate high-affinity binders, they carry an increased risk of immunogenicity, which may lead to premature CAR T cell clearance and treatment failure. Clinical studies have documented both humeral and cellular immune responses against murine- or camelid-derived CARs targeting antigens such as CD19, CD20, BCMA, and CAIX^27–32^. This immunogenicity may partly underly the limited efficacy of MGA271-based CARs observed in the STRlvE-02 and^16^ BrainChild-03 trials^6^•^15^. Although humanization strategies can reduce this risk, they may also compromise antibody stability or antigen binding due to the retention of non-human residues within CDRs and critical framework elements. In contrast, scFv phage display offers a robust, non-hybridoma approach to generate fully human antibodies. A key advantage of this method is the inherent selection of scFv stability during enrichment. Additionally, libraries derived from human B cell repertoires minimize immunogenic potential. This strategy has successfully yielded therapeutic antibodies including Avelumab (anti-PD-L1)^33^, Adalimumab (anti-TNF-a)^34^, and Necitumumab (anti-EGFR)^35^. To our knowledge, the present study represents the first report of fully human B7-H3-targeted CARs.

As with any solid tumor antigen, the potential for on-target off-tumor toxicity remains a concern. B7-H3 is expressed at low levels in certain normal tissues, including gastric epithelium, liver, and adrenal gland^2^ ^3^ Clinical studies to date have reported manageable toxicity with B7-H3 CART therapies^6^•^15^•^16^. Interestingly, Y111, selected for its strong binding to human B7-H3, also exhibits higher affinity for murine B7-H3 than benchmark CARs (Figure S3), enabling preliminary toxicity evaluation in immunodeficient mouse models. Body weights were unaltered, and no overt toxicity was observed (Figure S4). However, Y111’s affinity for mouse B7-H3 is approximately 17-fold lower than for the human antigen (Figure S3C), highlighting the need for careful toxicity monitoring as this CAR advances toward clinical evaluation.

The mechanisms underlying Y111’s markedly superior in vivo antitumor activity remain to be fully elucidated. Although binding affinity can influence CAR efficacy, no direct correlation was observed among the B7-H3 scFvs tested. Epitope location is another critical determinant of CAR function^19^•^20^, and Y111 may target a site that is particularly favorable for T cell activation. Additionally, Y111 may benefit from enhanced structural stability and proper folding, intrinsic to the phage display selection process. In contrast, benchmark CARs derived from murine binders, such as 376.96 and MGA271, require artificial VH-VL pairing to form scFvs, which can lead to reduced stability, impaired antigen binding, or unwanted tonic signaling. Notably, Y111 binds the 4Ig B7-H3 extracellular domain predominantly in a 1:1 monomeric fashion, potentially minimizing CAR clustering and tonic signaling caused by soluble B7-H3 in circulation.

In summary, this study reports the first fully human B7-H3-specific CART cells developed via phage display, yielding functionally stable and potentially less immunogenic binders. Among the candidates, Y111 CAR demonstrated superior and durable antitumor activity across multiple solid tumor models, outperforming existing clinical benchmark CARs. These results highlight the critical importance of binder selection in CAR T cell efficacy and underscore the potential of fully human antibodies to improve therapeutic outcomes in solid tumors. Further evaluation, including detailed epitope mapping and comprehensive safety profiling, will be essential as Y111 advances toward clinical development.

## MATERIALS AND METHODS

### Cell lines

HEK293T, Panc1, HPAC, and MIA PaCa-2 were obtained from the American Type Culture Collection (Rockville, MD). Nalm6 and NBEB were a kind gift from Dr. Javed Khan (NCI/NIH), IMRS and LAN5 were a kind gift from Dr. Mitchell Ho (NCI/NIH), and GBM01 were a kind gift from Dr. Anandani Nellan (NCI/NIH). Nalm6 cells were cultured in RPMl-1640 medium supplemented with 10% heat-inactivated fetal bovine serum (FBS) and 2% mM GlutaMAX. HEK293T cells were cultured in lscove’s Modified Dulbecco’s Medium supplemented with 10% heat-inactivated fetal bovine serum (FBS) and 2 mM GlutaMAX. All other cell lines were cultured in Dulbecco’s Modified Eagle’s Medium supplemented with 10% heat-inactivated fetal bovine serum (FBS) and 2 mM GlutaMAX. Cells were maintained in a 37°C incubator with a humidified environment containing 5% carbon dioxide (CO2). Mycoplasma testing was performed routinely on cell culture supernatant samples using MycoAlert Mycoplasma Detection Kit (Lonza, Rockland, ME, Catalog no. LT07-418) and all cell lines were confirmed to be negative. The identity of all cell lines was validated by Short Tandem Repeat (STR) Profiling, performed by the University of Arizona Genetics Core.

### Construction of the human naïve phage scFv library

B7-H3-specific binders were identified from a human naïve single-chain variable fragment (scFv) phage display library constructed in-house. Variable heavy (VH) regions derived from lgG and lgM, as well as kappa and lambda variable light (VL) regions, were amplified from RNA isolated from the bone marrow or peripheral blood mononuclear cells (PBMCs) of 58 healthy donors (bone marrow, n=50; PBMCs, n=8). VH and VL fragments were joined via overlapping PCR using a flexible Gly-Ser [(GS_4_h] linker, and Sfil restriction sites were incorporated at the 5’ and 3’ ends. The resulting scFv-encoding sequences were cloned into the Sfil site of the pADL1Ob vector (Antibody Design Laboratory). The pADL1Ob vector had been modified to include an OmpA signal peptide (MKKTAIAIAVALAGFATVAQA) upstream of the scFv insert, and tandem 6xHis and FLAG tags downstream. Recombinant pADL10b-scFv constructs were electroporated into E. coli TG1 competent cells (LGC Biosearch Technologies). Transformed TG1 cells were plated on Bioassay dishes (Sigma) to collect the library and on 10-cm plates to assess transformation efficiency. The final library consisted of four sub-libraries - lgG VH-kappa, lgG VH-lambda, lgM VH-kappa, and lgM VH-lambda - and contained approximately 8 x 10^10^ independent scFv clones.

### Selection of B7-H3-specific scFv binders

To identify binders recognizing both human and mouse B7-H3, 1 x 10^12^ plaque-forming units (PFU) of phages were preabsorbed with AP-biotin fusion protein (generated in-house) and Dynabeads MyOne Streptavidin T1 beads (Thermo Fisher). The precleared phages were then panned against 5 µg of biotinylated B7-H3-AP fusion protein. Streptavidin beads were used to capture bound phages, which were subsequently washed five times with PBS containing 0.05% Tween-20 (PBST). Bound phages were eluted and amplified in exponentially growing TG1 cells in the presence of M13KO7 helper phage (Antibody Design Laboratory). Two additional rounds of panning were performed under increasingly stringent conditions: round 2 used 2 µg of biotinylated B7-H3-AP and 15 PBST washes, and round 3 used 1 µg of antigen and 20 washes. Following three rounds of selection, 400 individual clones were rescued with helper phage and screened by phage ELISA for binding to human and mouse B7-H3 proteins. Positive scFv clones were further validated in cell-based binding assays.

### scFv production and purification

Selected scFv clones were expressed in E. coli HB2151 (Life Science Market) harboring the pADL1Ob vector. Single bacterial colonies were inoculated into 2xYT medium (Sigma) supplemented with 50 µg/ml ampicillin and 0.2% glucose (Sigma) and cultured at 37 °C with shaking until the optical density at 600 nm (OD_600_) reached 0.9. Protein expression was induced by the addition of 1 mM IPTG (Gold Biotechnology), followed by incubation at 30 °C overnight with shaking. Cells were lysed by treatment with polymyxin B (10,000 U/ml; Sigma), and the soluble scFv fraction was recovered from the clarified lysate by centrifugation. scFv proteins were purified using Ni^2^+-NTA affinity chromatography. The column was washed with PBS containing 0.5 M NaCl and 20 mM imidazole (Sigma) to remove non-specifically bound proteins, and scFvs were eluted with PBS containing 0.5 M NaCl and 200 mM imidazole. The eluted scFv proteins were further purified by size-exclusion chromatography on a Superdex 200 Increase 10/300 GL column (Cytiva) to remove dimeric species. The monomeric scFv fractions were concentrated and dialyzed against PBS using 3 kDa MWCO dialysis cassettes (Thermo Scientific) to obtain purified scFv proteins for downstream applications.

### Generation of 87-H3 CAR vectors

The SFG retroviral backbone utilized to generate B7-H3 CAR vectors for this study has been previously described^36^. The constructs encoding B7-H3 CARs derived from all binders described in the study (Y868, Y111, Y117, MGA271, 376.96, and 812) were codon optimized and synthesized by Integrated DNA Technologies (Coralville, IA). The encoding sequences were then subcloned into the retroviral backbone by Gibson Assembly using Gibson Assembly Cloning Kit (NEB). The sequences of all B7-H3 CAR constructs were validated by sequencing (Quintara Bio). Retrovirus-containing supernatant for transduction of T cells was produced as described previously^37^. Briefly, 293T cells were co-transfected with the CAR-encoding plasmid, the RD114 envelope plasmid, and the MomLV gag-pol encoding pEQ-Pam3(-E) plasmid using GeneJuice Transfection Reagent (Sigma-Aldrich, USA). Supernatant was collected 24, 48, and 72 hours after transfection, filtered using 0.45 µm Millex™-HV Filter Unit (Millipore Sigma), and stored at - 80 °C until use.

### CAR T cell production

Generation of CAR T cells have been described previously^38^. Briefly, peripheral blood mononuclear cells (PBMCs) were isolated from healthy donor blood by density gradient separation using Lymphoprep (Stemcell Technologies). Donor blood samples were obtained through the Research Donor Program (RDP) at the National Cancer Institute - Frederick Occupational Health Services Clinic from volunteers under a study protocol approved by the National Institutes of Health Institutional Review Board. 1×10^6^ PBMCs were seeded in each well of untreated 24-well plates pre-coated with anti-CD3 and anti-CD28 antibodies to selectively activate T cells. After overnight incubation, IL-2 was added to the culture medium at the final concentration of 100 IU/ml. 48 hours after incubation in the antibody-coated wells, T cells were harvested and transduced to express CARs using retroviral virus supernatant as described previously^38^.

### Generation of luciferase-labeled tumor cells

Tumor cells were transduced with a retroviral vector encoding a GFP and firefly luciferase fusion protein (GL) to stably express these proteins to facilitate the detection of tumor cells by flow cytometry, luminescence-capable spectrophotometer, and in vivo bioluminescence imaging systems. The retroviral vector used to engineer tumor cells have been described in a prior publication^39^. Virus supernatant was obtained by collecting conditioned medium from cultures of a producer cell line engineered to produce the GL encoding virus particles. Supernatant was used to transduce tumor cell lines using the protocol used to transduce T cells. Transduction efficiency was determined based on the detection of GFP expressed by the engineered cancer cells by flow cytometry. All cell lines were sorted to exclude GFP-negative cells prior to use in the experiments described in this study.

### B7-H3 Gene Targeting in NBEB and Panc-1 Cells Using CRISPR-Cas9

The B7-H3 KO construct was generated using two guide RNA sequences: CD276-guide-1 (5’-CACCGTGGCACAGCTCAACCTCATC-3’) and CD276-guide-2 (5’-AAACGATGAGGTTGAGCTGTGCCAC-3’) (Integrated DNA Technologies). Oligonucleotides were phosphorylated with T4 polynucleotide kinase (PNK; NEB, M0201S), annealed at 95°C for 5 min, gradually cooled to room temperature, and ligated into BsmBl-digested lentiCRISPR v2 vector (Addgene plasmid #52961) using Quick Ligase (NEB, M2200S). Lentiviral particles were produced by transfecting Lenti-X 293T cells (Takara, 632180) with the gRNA-expressing lentiCRISPR v2 plasmid, pMD2.G (Addgene, 12259), and psPAX2 (Addgene, 12260) at a 3:2:5 ratio using Lipofectamine 2000 (lnvitrogen, 11668019) according to the manufacturer’s instructions. Viral supernatants were concentrated using Lenti-X Concentrator (Takara, 631232) and used to transduce target NBEB and Panc-1 cells. Forty-eight to seventy-two hours post-transduction, puromycin selection was applied to enrich for transduced cells. One week after selection, cells were stained with rabbit anti-B7-H3 antibody (clone EPNCIR122, Abeam, ab134161) and sorted by flow cytometry to isolate B7-H3-negative populations. Parental B7-H3-positive cells were used to establish gating, and cells were subjected to a second round of sorting to ensure a homogeneous B7-H3 knockout population.

### Flow cytometry

Labeling of cell surface proteins for flow cytometry was performed by standard staining protocol - cells were incubated with the manufacturer-recommended amounts of antibody (∼5 µL) for 20 min at 4°C and washed with 1 x Dulbecco’s PBS (DPBS; Corning, catalog no. 21-031-CV). Intracellular cytokine staining (ICS) was performed according to the instructions provided by the manufacturer with the ICS reagents/antibodies. All samples were analyzed using either an LSR II SORP (Becton Dickinson) or a MACSQuant 16 (Miltenyi) flow cytometer. CAR expression by **T** cells was measured 3-4 days post-transduction. CARs expressed on the surface of T cells were detected using either an antibody or recombinant antigen. For the antibody mediated detection, CAR transduced **T** cells were incubated with goat anti-human F(ab’)2 antibody conjugated with Alexa Fluor 647 (Jackson lmmunoResearch Laboratories, West Grove, PA, catalog no. 109-606-097) before flow analysis. For the antigen method of CAR detection, **T** cells were first incubated with biotin-tagged extracellular domain of human B7-H3, followed by incubation with APC-conjugated Streptavidin (BioLegend). The following antibodies used for T cell phenotyping and ICS were all obtained from BioLegend: CD3-PerCPCy5.5, CD4-AF700, CD8-PB, CCR7-PE/Cy7, CD45RA-APC, CD69-APC Cy7, LAG3-PE/Cy7, PD1-APC, TIM3-FITC, Granzyme-B-FITC, Perforin-APC, CD107a-PE/Dazzle. **T** cells were fixed and permeabilized using Fixation Buffer and Intracellular Staining Permeabilization Wash Buffer, respectively, according to the ICS protocol from BioLegend. B7-H3 expression by cancer cell lines used in the study was measured using B7-H3-PE antibody (Miltenyi Biotec, catalog no. 130-120-781). To compare B7-H3 detection by Y868, Y111, and Y1117, purified scFvs were incubated at indicated concentrations with cancer cells, followed by AF-647 (Panc1 and IMR5) or PE (293T) conjugated anti-human F(ab’)2 secondary antibodies. Flow cytometry data were analyzed using FlowJo version 10.9.0 or version 11 (Becton Dickinson).

### Luciferase-based killing assay

CAR T cell anti-tumor activity in vitro was measured co-culture experiments in which T cells were incubated with GL-labeled tumor cell lines. Cell lines naturally lacking B7-H3 expression or B7-H3 knock-out tumor cell lines, also transduced to express GL were used as negative controls. 2×10^7^ tumor cell lines were seeded in 96-well black flat-bottom tissue culture-treated plates (Corning) in 100 µL of T cell medium. After 2 hour incubation at 37°C, un-transduced (UTD) or CAR-transduced T cells were resuspended at various cell concentrations in cytokine-free T cell medium and 100 µL of T cell suspension was added to the tumor cells resulting in effector to target ratios (E:T) of 5:1, 2.5:1, 1.25:1, 0.625:1, and 0.25:1. All conditions were setup in triplicates. After 48-hour incubation at 37°C, D-Luciferin (GoldBio, catalog no. LUCK-1G) was added to each well at the final concentration of 5 µg/ml. Immediately after addition of D-Luciferin, bioluminescence produced by the remaining tumor cells was measured using a CLARIOstar microplate reader (BMG LABTECH, Cary, NC). Background luminescence signals from wells containing tumor cells without D-Luciferin addition was also captured and subtracted from the readouts obtained from sample wells. Using the background-subtracted luminescence values, percent (%) cytotoxicity was calculated using the following formula: [(Tumor-only luminescence - Treated-condition luminescence)/ Tumor-only luminescence] x 100. In some experiments, T cells were collected for flow cytometry to measure intracellular cytokines or degranulation markers, after obtaining the luminescence data. Briefly, T cells in the wells were re-suspended in the culture medium by repeated gentle pipetting and transferred to flow cytometry sample tubes (Falcon-A Corning Brand, catalog no. 352052) for labeling according to description provided in the flow cytometry section. Conditioned media from some experiments was also collected carefully without disturbing the cells and stored at −80°C to measure the production of cytokines by T cells during co-culture with tumor cells.

### Enzyme-Linked lmmunosorbent Assay (ELISA) for Cytokine quantification

Cytokine production by T cells cocultured with tumor cells was quantified using ELISAs. Conditioned media collected from luciferase-based cytotoxicity assays were analyzed using Granzyme B, IFN-y, and IL-2 ELISA kits, following the manufacturers’ protocols. The E:T ratio used for ELISA experiments was 2.5:1. Absorbance was measured using a CLARIOstar microplate reader (BMG Labtech).

### Epitope mapping

To map antibody binding regions, ELISA plates were coated overnight at 4°C with purified B7-H3 protein or PD-L1/B7-H3 chimeric fusion proteins. Wells were blocked with 2% milk in PBS, and antibodies were added in triplicate at final concentrations of 0.00128, 0.0064, 0.032, 0.16, 0.8, 4, 20, and 100 nM. Plates were incubated at 37°C for 2 h, washed with PBS containing 0.05% Tween-20 (PBST), and then incubated for 1 h at 37°C with horseradish peroxidase (HRP)-conjugated secondary antibodies. Donkey anti-human Fc-HRP (Jackson lmmunoResearch Laboratories) was used to detect Y111 and MGA271 lgGs, while goat anti-mouse lgG-HRP (Southern Biotech) was used for 376.96 lgG detection. After washing with PBST, bound antibodies were visualized using ABTS substrate (Sigma), and absorbance was recorded on the CLARIOstar plate reader.

### Animal studies

All animal studies were performed using 6- to 8-week-old female NRG mice (NOD.Cg-Rag1tm1Mom ll2rgtm1Wjl/SzJ, Jackson Laboratory strain no. 007799) under protocols approved by the National Cancer Institute Institutional Animal Care and Use Committee. Xenograft tumors of pancreatic ductal adenocarcinoma, neuroblastoma, glioblastoma (GBM), and rhabdomyosarcoma were established using respective wildtype or GL-labeled lines, doses, and injection routes as described or illustrated in the manuscript text, figures, or figure legends. Tumor burden was measured using calipers for subcutaneous and rhabdomyosarcoma tumor models.

In vivo bioluminescence imaging (BU) using IVIS Spectrum or IVIS Lumina Ill optical scanners (PerkinElmer) were used to monitor tumor progression or regression in other tumor models, as described or illustrated in the manuscript text and figures. For tumors measured using calipers, tumor volumes were calculated using the formula: length x width x width/2. For BU-mediated tumor measurements, anesthetized mice were imaged once a week or at indicated time-points to capture tumor luminescence signal 10 minutes after intraperitoneally injecting 100 µL of 15 mg/ml D-Luciferin solution in 1X PBS. BU imaging data was imported into and analyzed using Aura version 4.0.8 or 5.0.1 (Spectral Instruments) or Living Image software, version 4.7.4 (PerkinElmer) to quantify the BU signal from tumors. T cells were injected intravenously via tail vein for all tumor models used in the study. T cell doses for each tumor study are indicated in the main text, figures, or figure legends. Mice were humanely euthanized when the tumor volume reached the protocol limit of 4000 mm^3^ for subcutaneous tumors. In the mice with BU-monitored tumors, symptoms of tumor-caused illnesses (e.g., hunched posture, ruffled coat, lack of mobility, head tilt, and weight loss) or high tumor burden based on bioluminescence imaging were used as endpoint criteria to humanely euthanize the mice. For the tumor-rechallenge arm of the HPAC study, mice free of initially implanted orthotopic tumor were subcutaneously injected in the lower right abdomen with 4×10^6^ HPAC-GL cells. Age matched naïve (no previous administration of tumor or T cells) female NRG mice were also injected with tumor cells at the same time to serve as controls. All subsequent tumor measurements were performed using BU for both the initial tumor in the pancreas and the subcutaneous tumor at the rechallenge site.

### Statistical analysis

All data in this manuscript are reported as average± standard error of mean (SEM). Initial data recording and/or preliminary analysis/organization was performed in Microsoft Excel. Final analysis, generation of data graphs, and statistical analyses were all performed in GraphPad Prism. Analyses performed to determine statistical significance are indicated in the figure legends. P value ≤0.05 was considered statistically significant. ns = no significant difference (P>0.05), * = P≤0.05, ** = P≤0.01, *** = P≤0.001, **** = P≤0.0001.

## RESOURCE AVAILABILITY

### Lead contact

Further information and requests for resources and reagents should be directed to and will be fulfilled by the lead contact, Brad St. Croix ixb@nih.g

### Materials availability

Reagents generated in this study, including B7-H3-CAR T vectors, will be made available upon request. Some materials may require requests to collaborators and/or agreements with various entities. Materials that can be shared will be released via a Material Transfer Agreement.

### Data and code availability

This paper analyzes existing, publicly available data. The cancer mRNA data are available from the Depmap portal website: https://depmap.org/portal/, 2204 release. This paper does not report original code. Any additional information required to reanalyze the data reported in this paper will be shared by the lead contact upon request.

## ACKNOWLEDGMENTS

This research was supported by the Intramural Research Program of the National Institutes of Health (NIH) and a NCI CCR FLEX Program Synergy Award (B.S.C). We thank the NCI CCR-Frederick Flow Cytometry Core Facility for assistance in cell sorting and analysis. We thank the Small Animal Imaging Program (SAIP) for facilitating bioluminescence imaging and veterinary staff for monitoring the health of mice. We also thank Dr. Ann Leen from Baylor College of Medicine for the eGFP-FFluc vector producer cell line. The anti-B7H3 scFv antibodies presented in the present study are the subject of pending patent applications assigned to the NIH and are available for license in certain fields of use to qualified candidates. Please contact the corresponding author Dr. St Croix (NCI) at if you are interested in pursuing a license. The contributions of the NIH authors were made as part of their official duties as NIH federal employees, are in compliance with agency policy requirements, and are considered Works of the United States Government. However, the findings and conclusions presented in this paper are those of the authors and do not necessarily reflect the views of the NIH or the U.S. Department of Health and Human Services.

## AUTHOR CONTRIBUTIONS

Conceptualization, P.B., Y.F., and B.S.C.; Investigation, P.B., Y.F., M.P., M.T., K-S.H., J.L., G.Y., L.Y., S.S., M.B.H., K.M., N.B., J.T., and R.D.M.; Writing - Original Draft, P.B. and B.S.C.; Writing - Review & Editing, all authors; Funding Acquisition, B.S.C., J.K., AN., R.N.K. and B.S.C.; Supervision, J.C.C., J.K., A.N., R.N.K and B.S.C.

## DECLARATION OF INTERSTS

P.B., Y.F. and B.S.C. are NIH inventors of a provisional patent application (International application number PCT/US2024/037349) filed by the United States of America, as represented by the Secretary, Department of Health and Human Services, for the use of fully human chimeric antigen receptors against CD276 for the treatment of solid tumors. The remaining authors declare no competing interests.

## References

1. Cappell, K.M., and Kochenderfer, J.N. (2023). Long-term outcomes following CART cell therapy: what we know so far. Nat Rev Clin Oneel 20, 359–371. 10.1038/s41571-023-00754-1.

2. Seaman, S., Zhu, Z., Saha, S., Zhang, X.M., Yang, M.Y., Hilton, M.B., Morris, K., Szot, C., Morris, H., Swing, D.A, et al. (2017). Eradication of Tumors through Simultaneous Ablation of CD276/B7-H3-Positive Tumor Cells and Tumor Vasculature. Cancer Cell 31, 501–515 e508. 10.1016/j.ccell.2017.03.005.

3. Du, H., Hirabayashi, K., Ahn, S., Kren, N.P., Montgomery, S.A, Wang, X., Tiruthani, K., Mirlekar, B., Michaud, D., Greene, K., et al. (2019). Antitumor Responses in the Absence of Toxicity in Solid Tumors by Targeting B7-H3 via Chimeric Antigen Receptor T Cells. Cancer Cell 35, 221–237 e228. 10.1016/j.ccell.2019.01.002.

4. Majzner, R.G., Theruvath, J.L., Nellan, A, Heitzeneder, S., Cui, Y., Mount, C.W., Rietberg, S.P., Linde, M.H., Xu, P., Rota, C., et al. (2019). CART Cells Targeting B7-H3, a Pan-Cancer Antigen, Demonstrate Potent Preclinical Activity Against Pediatric Solid Tumors and Brain Tumors. Clin Cancer Res 25, 2560–2574. 10.1158/1078-0432.CCR-18-0432.

5. Epperly, R., Gottschalk, S., and DeRenzo, C. (2024). CART cells redirected to B7-H3 for pediatric solid tumors: Current status and future perspectives. EJC Paediatr Oneel 3. 10.1016/j.ejcped.2024.100160.

6. Pinto, N., Albert, C.M., Taylor, M.R., Ullom, H.B., Wilson, AL., Huang, W., Wendler, J., Pattabhi, S., Seidel, K., Brown, C., et al. (2024). STRlvE-02: A First-in-Human Phase I Study of Systemically Administered B7-H3 Chimeric Antigen Receptor T Cells for Patients With Relapsed/Refractory Solid Tumors. J Clin Oneel, JCO2302229. 10.1200/JC0.23.02229.

7. Khan, AN., Chowdhury, A, Karulkar, A, Jaiswal, AK., Banik, A, Asija, S., and Purwar, R. (2022). lmmunogenicity of CAR-T Cell Therapeutics: Evidence, Mechanism and Mitigation. Front lmmunol 13, 886546. 10.3389/fimmu.2022.886546.

8. Wutti-ln, Y., Sujjitjoon, J., Sawasdee, N., Panya, A, Kongkla, K., Yuti, P., Yongpitakwattana, P., Thepmalee, C., Junking, M., Chieochansin, T., et al. (2021). Development of a Novel Anti-CD19 CAR Containing a Fully Human scFv and Three Costimulatory Domains. Front Oncol 11, 802876. 10.3389/fonc.2021.802876.

9. Abbott, R.C., Verdon, D.J., Gracey, F.M., Hughes-Parry, H.E., lliopoulos, M., Watson, K.A, Mulazzani, M., Luong, K., D’Arcy, C., Sullivan, L.C., et al. (2021). Novel high-affinity EGFRvlll-specific chimeric antigen receptor T cells effectively eliminate human glioblastoma. Clin Transl Immunology 10, e1283. 10.1002/cti2.1283.

10. Imai, K., Wilson, B.S., Bigotti, A, Natali, P.G., and Ferrone, S. (1982). A 94,000-dalton glycoprotein expressed by human melanoma and carcinoma cells. J Natl Cancer Inst 68, 761–769.

11. Loo, D., Alderson, R.F., Chen, F.Z., Huang, L., Zhang, W., Gorlatov, S., Burke, S., Ciccarone, V., Li, H., Yang, Y., et al. (2012). Development of an Fe-enhanced anti-B7-H3 monoclonal antibody with potent antitumor activity. Clin Cancer Res 18, 3834–3845. 10.1158/1078-0432.CCR-12-0715.

12. Li, D., Wang, R., Liang, T., Ren, H., Park, C., Tai, C.H., Ni, W., Zhou, J., Mackay, S., Edmondson, E., et al. (2023). Camel nanobody-based B7-H3 CAR-T cells show high efficacy against large solid tumours. Nat Commun 14, 5920. 10.1038/s41467-023-41631-w.

13. Birley, K., Leboreiro-Babe, C., Rota, E.M., Buschhaus, M., Gavriil, A, Vitali, A, Alonso-Ferrero, M., Hopwood, L., Parienti, L., Ferry, G., et al. (2022). A novel anti-B7-H3 chimeric antigen receptor from a single-chain antibody library for immunotherapy of solid cancers. Mol Ther Oncolytics 26, 429–443. 10.1016/j.omto.2022.08.008.

14. Majzner, R.G., Rietberg, S.P., Sotillo, E., Dong, R., Vachharajani, V.T., Labanieh, L., Myklebust, J.H., Kadapakkam, M., Weber, E.W., Tousley, AM., et al. (2020). Tuning the Antigen Density Requirement for CAR T-cell Activity. Cancer Discov 10, 702–723. 10.1158/2159-8290.CD-19-0945.

15. Vitanza, N.A., Ronsley, R., Choe, M., Seidel, K., Huang, W., Rawlings-Rhea, S.D., Beam, M., Steinmetzer, L., Wilson, A.L., Brown, C., et al. (2025). lntracerebroventricular B7-H3-targeting CART cells for diffuse intrinsic pontine glioma: a phase 1 trial. Nat Med 31, 861–868. 10.1038/s41591-024-03451-3.

16. Vitanza, N.A., Wilson, AL., Huang, W., Seidel, K., Brown, C., Gustafson, J.A., Yokoyama, J.K., Johnson, A.J., Baxter, B.A., Koning, R.W., et al. (2023). lntraventricular B7-H3 CAR T Cells for Diffuse Intrinsic Pontine Glioma: Preliminary First-in-Human Bioactivity and Safety. Cancer Discov 13, 114–131. 10.1158/2159-8290.CD-22-0750.

17. Li, G., Stocksdale, B.R., Song, K.-W., Mahdi, J., Tanner, K., Bertrand, S., Iv, M., Threlkeld, Z., Ramakrishna, S., Sahaf, B., et al. (2025). A phase 1 study of B7H3 CAR-T cells administered intracranially in recurrent glioblastoma. J Clin Oncol 43 (16), p2018.

18. Tang, X., Wang, Y., Huang, J., Zhang, Z., Liu, F., Xu, J., Guo, G., Wang, W., Tong, A., and Zhou, L. (2021). Administration of B7-H3 targeted chimeric antigen receptor-T cells induce regression of glioblastoma. Signal Transduct Target Ther 6, 125. 10.1038/s41392-021-00505-7.

19. James, S.E., Greenberg, P.O., Jensen, M.C., Lin, Y., Wang, J., Till, B.G., Raubitschek, A.A., Forman, S.J., and Press, O.W. (2008). Antigen sensitivity of CD22-specific chimeric TCR is modulated by target epitope distance from the cell membrane. J lmmunol 180, 7028–7038. 10.4049/jimmunol.180.10.7028.

20. Liu, X., Onda, M., Watson, N., Hassan, R., Ho, M., Bera, T.K., Wei, J., Chakraborty, A., Beers, R., Zhou, Q., et al. (2022). Highly active CART cells that bind to a juxtamembrane region of mesothelin and are not blocked by shed mesothelin. Proc Natl Acad Sci U S A 119, e2202439119. 10.1073/pnas.2202439119.

21. Chang, Z.L., Lorenzini, M.H., Chen, X., Tran, U., Bangayan, N.J., and Chen, Y.Y. (2018). Rewiring T-cell responses to soluble factors with chimeric antigen receptors. Nat Chem Biol 14, 317–324. 10.1038/nchembio.2565.

22. Zhou, Y.H., Chen, Y.J., Ma, Z.Y., Xu, L., Wang, Q., Zhang, G.B., Xie, F., Ge, Y., Wang, X.F., and Zhang, X.G. (2007). 4IgB7-H3 is the major isoform expressed on immunocytes as well as malignant cells. Tissue Antigens 70, 96–104. 10.1111/j.1399-0039.2007.00853.x.

23. Steinberger, P., Majdic, 0., Derdak, S.V., Pfistershammer, K., Kirchberger, S., Klauser, C., Zlabinger, G., Pickl, W.F., Stocki, J., and Knapp, W. (2004). Molecular characterization of human 4Ig-B7-H3, a member of the B7 family with four lg-like domains. J lmmunol 172, 2352–2359. 10.4049/jimmunol.172.4.2352.

24. Azuma, T., Sato, Y., Ohno, T., Azuma, M., and Kume, H. (2020). Serum soluble B7-H3 is a prognostic marker for patients with non-muscle-invasive bladder cancer. PLoS One 15, e0243379. 10.1371/journal.pone.0243379.

25. Zhang, G., Xu, Y., Lu, X., Huang, H., Zhou, Y., Lu, B., and Zhang, X. (2009). Diagnosis value of serum B7-H3 expression in non-small cell lung cancer. Lung Cancer 66, 245–249. 10.1016/j.lungcan.2009.01.017.

26. Avci, 0., Cavdar, E., lriagac, Y., Karaboyun, K., Celikkol, A., Ozcaglayan, T.I.K., Oznur, M., Gurdal, S.O., and Seber, E.S. (2022). Soluble B7H3 level in breast cancer and its relationship with clinicopathological variables and T cell infiltration. Contemp Oncol (Pozn) 26, 27–31. 10.5114/wo.2022.113502.

27. Lamers, C.H., Willemsen, R., van Elzakker, P., van Steenbergen-Langeveld, S., Broertjes, M., Oosterwijk-Wakka, J., Oosterwijk, E., Sleijfer, S., Debets, R., and Gratama, J.W. (2011). Immune responses to transgene and retroviral vector in patients treated with ex vivo-engineered T cells. Blood 117, 72–82. 10.1182/blood-2010-07-294520.

28. Bartolo-lbars, A., Aran, A., Johansson, AM., Ortiz-Maldonado, V., Klein-Gonzalez, N., Alonso-Saladrigues, A., James, E., Urbano-lspizua, A., Rives, S., Delgado, J., et al. (2025). T-cell responses against CD19-targeted CART cells varnimcabtagene autoleucel (ARl-0001): implications for immune response and therapy outcomes. J lmmunother Cancer 13. 10.1136/jitc-2024-010792.

29. Lam, N., Trinklein, N.D., Buelow, B., Patterson, G.H., Ojha, N., and Kochenderfer, J.N. (2020). Anti-BCMA chimeric antigen receptors with fully human heavy-chain-only antigen recognition domains. Nat Commun 11, 283. 10.1038/s41467-019-14119-9.

30. Lamers, C.H., Klaver, Y., Gratama, J.W., Sleijfer, S., and Debets, R. (2016). Treatment of metastatic renal cell carcinoma (mRCC) with CAIX CAR-engineered T-cells-a completed study overview. Biochem Soc Trans 44, 951–959. 10.1042/BST20160037.

31. Wagner, D.L., Fritsche, E., Pulsipher, M.A., Ahmed, N., Hamieh, M., Hegde, M., Ruella, M., Savoldo, B., Shah, N.N., Turtle, C.J., et al. (2021). lmmunogenicity of CART cells in cancer therapy. Nat Rev Clin Oncol 18, 379–393. 10.1038/s41571-021-00476-2.

32. Till, B.G., Jensen, M.C., Wang, J., Chen, E.Y., Wood, B.L., Greisman, H.A., Qian, X., James, S.E., Raubitschek, A., Forman, S.J., et al. (2008). Adoptive immunotherapy for indolent non-Hodgkin lymphoma and mantle cell lymphoma using genetically modified autologous CD20-specific T cells. Blood 112, 2261–2271. 10.1182/blood-2007-12-128843.

33. Kim, E.S. (2017). Avelumab: First Global Approval. Drugs 77, 929–937. 10.1007/s40265-017-0749-6.

34. Bain, B., and Brazil, M. (2003). Adalimumab. Nat Rev Drug Discov 2, 693–694. 10.1038/nrd1182.

35. Garnock-Jones, K.P. (2016). Necitumumab: First Global Approval. Drugs 76, 283–289. 10.1007/s40265-015-0537-0.

36. Riviere, I., Brose, K., and Mulligan, R.C. (1995). Effects of retroviral vector design on expression of human adenosine deaminase in murine bone marrow transplant recipients engrafted with genetically modified cells. Proc Natl Acad Sci U S A 92, 6733–6737. 10.1073/pnas.92.15.6733.

37. Leen, A.M., Sukumaran, S., Watanabe, N., Mohammed, S., Keirnan, J., Yanagisawa, R., Anurathapan, U., Rendon, D., Heslop, H.E., Rooney, C.M., et al. (2014). Reversal of tumor immune inhibition using a chimeric cytokine receptor. Mol Ther 22, 1211–1220. 10.1038/mt.2014.47.

38. Torres Chavez, A.G., McKenna, M.K., Gupta, A., Daga, N., Vera, J., Leen, A.M., and Bajgain, P. (2025). IL-7 armed binary CAR T cell strategy to augment potency against solid tumors. Front lmmunol 16, 1618404. 10.3389/fimmu.2025.1618404.

39. Vera, J., Savoldo, B., Vigouroux, S., Biagi, E., Pule, M., Rossig, C., Wu, J., Heslop, H.E., Rooney, C.M., Brenner, M.K., and Dotti, G. (2006). T lymphocytes redirected against the kappa light chain of human immunoglobulin efficiently kill mature B lymphocyte-derived malignant cells. Blood 108, 3890–3897. 10.1182/blood-2006-04-017061.

