## Supplementary Data for "Engineering Functionality Optimized fully human B7-H3 CAR T Cells for Enhanced Solid Tumor Therapy"

**A**

| scFv clone | Sequences |
|---|---|
| Y868 | **$V_H$ Sequence**<br>CDR1 CDR2 CDR3<br>QVQLVESGGGLVQPGGSLRLSCAASGFTFSSYAMSWVRQAPGKGLEWVSAISGSGGSTYYADSVKGRFTISRDNAKNSLYLQVNSLRAEDTALYYCVRDTYRFFDYWGQGTLVTVSS<br>**$V_L$ Sequence**<br>CDR1 CDR2 CDR3<br>QSVLTQPPSVSGAPGQRVTISCSGSTSNIGARYPVHWYQQFPGTVPKLLIYGNNNRPSGVPDRFSGSKSGTSASLVITGLQAGDEADYYCQSFDNSLSGVLFGGGTKVTVLG |
| Y111 | **$V_H$ Sequence**<br>CDR1 CDR2 CDR3<br>QVQLVQSGAEVKKPGASVKVSCKASGYTFTSYGISWVRQAPGQGLEWMGRIIPILGIANYAQKFQGRVTITADKSTSTAYMELSSLRSEDTAVYYCARSNPGGAFDIWGQGTMVTVSS<br>**$V_L$ Sequence**<br>CDR1 CDR2 CDR3<br>DIQLTQSPSTLSASVGDRVTITCRASQSVRSNLAWYHQKPGKAPKLLIYKASTLESGVPSRISGSGSGTEFTLTISSLQPDDFATYFCQQYAYYPTFGQGTRLEIKG |
| Y117 | **$V_H$ Sequence**<br>CDR1 CDR2 CDR3<br>QVQLVQSGAEVKQPGASVKVSCKASGYTFTDYAITWVRQAPGQGLEWVGWINTDTGNPTYAQAFTGRLVFSLDTSVSTAYLQISSLKAEDTAVYYCARVTPSGSLVHYDFWGQGTLVTVSS<br>**$V_L$ Sequence**<br>CDR1 CDR2 CDR3<br>QSVLTQPPSVSGAPGQRVTISCTGSRSNTGAGFDVHWYQQLPGTAPKLLIYGNNNRPSGVPDRFSGSRSGTSASLAINGLQAEDEADYYCQSYDYNLSGVIFGGGTKLTALG |

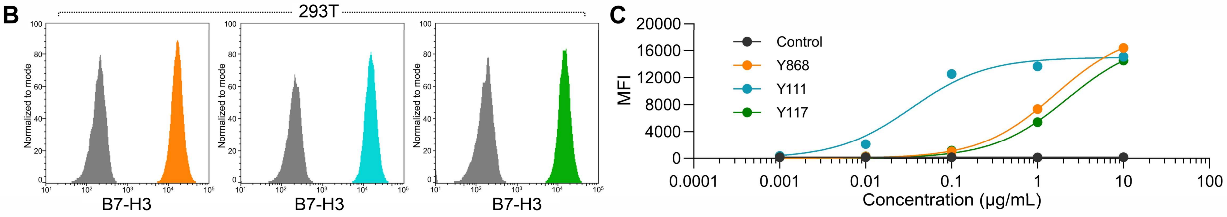


**Supplemental Figure 1. B7-H3 antibodies contain distinct variable domains and bind specifically to B7-H3.**
(A) Variable domain sequences of Y868, Y111 and Y117.
(B) Flow cytometry evaluating binding of Y868, Y111 and Y117 to 293T cells.
(C) Flow cytometry showing the dose-dependent binding of each of the scFvs. MFI: mean fluorescence Intensity.

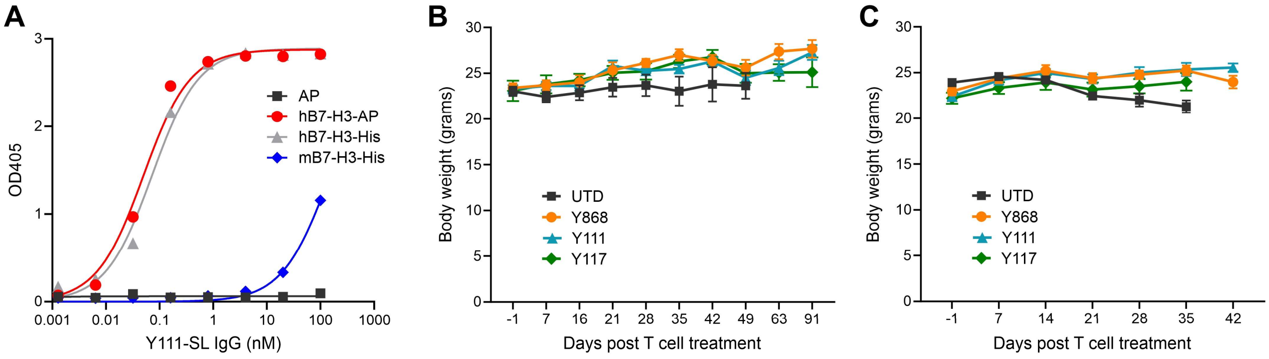


**Supplemental Figure 2. B7-H3 CAR T cells are well tolerated in vivo.**
(A) ELISA showing reactivity of Y111 full IgG to Alkaline phosphatase (AP) control protein, or the extracellular domain (ED) of human B7-H3 fused to AP (hB7-H3-AP) or 6xHIS (hB7-H3-His), or mouse b7-H3 ED fused to 6xHIS (mB7-H3-His).
(B) Body weight measurements following CAR T treatment for mice in the Panc1 study shown in Figure 4A-D of the main text.
(C) Body weight measurements following CAR T treatment for mice in the HPAC study shown in Figure 4E-G of the main text.

**A**

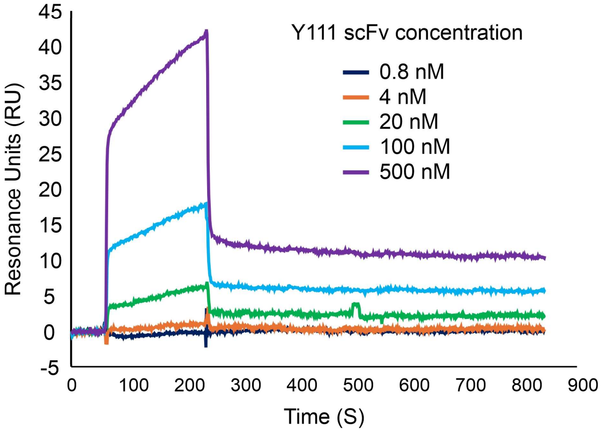


**B**

| | | human B7H3-ED | | | mouse B7H3-ED | | |
|---|---|---|---|---|---|---|---|
| | mAb | ka (1/Ms) | kd (1/s) | KD (nM) | ka (1/Ms) | kd (1/s) | KD (nM) |
| IgG | Y111 | $1.2 \times 10^{6}$ | $2.6 \times 10^{-2}$ | 21.1 | $1.4 \times 10^{5}$ | 0.05 | 366.2 |
| | MGA271 | $1.9 \times 10^{5}$ | $1.1 \times 10^{-3}$ | 5.7 | ND | ND | ND |
| | 376.96 | $5.3 \times 10^{4}$ | $1.1 \times 10^{-3}$ | 20.9 | $1.0 \times 10^{4}$ | 0.02 | 2040.1 |
| scFv | Y111 | $4.3 \times 10^{4}$ | $5.0 \times 10^{-4}$ | 11.8 | - | - | - |
| | MGA271 | $1.7 \times 10^{5}$ | $1.5 \times 10^{-2}$ | 91.0 | - | - | - |
| | 376.96 | $1.7 \times 10^{5}$ | $2.0 \times 10^{-2}$ | 116.9 | - | - | - |

ND: no detectable binding

**Supplemental Figure 3. Affinity measurements for Y111, MGA271, and 376.96 antibodies.** (A) Biacore sensogram showing the binding kinetics of Y111 scFv to immobilized B7-H3 extracellular domain (ED). Y111 exhibits rapid initial binding, followed by a second, concentration-dependent association phase. B7-H3 contains an internal repeat within its ED, and Y111 preferentially forms a 1:1 complex with B7-H3 (see Supplemental Fig. 6D). The weaker secondary association likely reflects binding of a second scFv at the unoccupied repeat in the 1:1 complex at high Y111 scFv concentrations. During the dissociation phase, the resonance units (RU) decrease rapidly, followed by a slower decline, consistent with rapid dissociation from the secondary, lower-affinity site and slower release from the primary binding site.

(B) Summary of binding affinities of Y111, MGA271, and 376.96 in either full IgG (bivalent) or scFv (monovalent) formats against B7-H3 ED. Y111 is unusual in that it exhibits higher affinity in monovalent scFv format, indicating that avidity of the antibody does not contribute substantially to its binding strength.

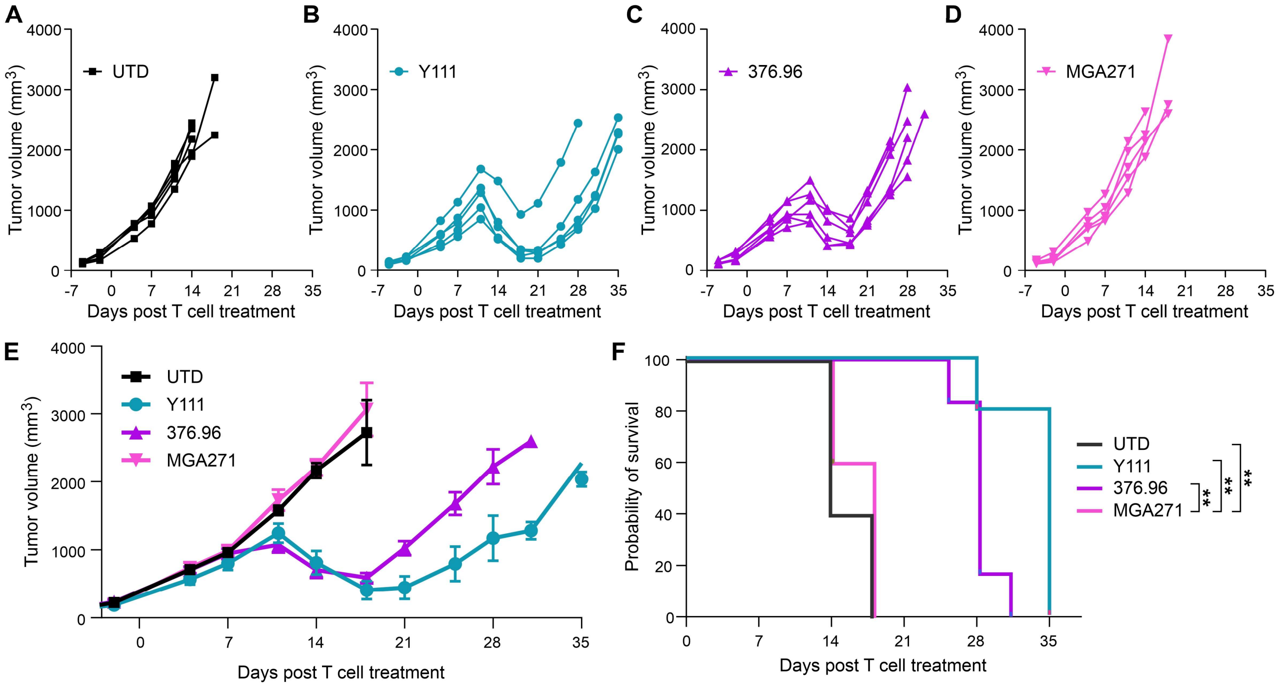


**Supplemental Figure 4. Y111 compares favorably with benchmark B7-H3 CARs in a rhabdomyosarcoma model.**

(A-D) Individual tumor growth curves of JR-1 rhabdomyosarcoma following intramuscular tumor implantation and subsequent i.v. administration of (A) UTD control, (B) Y111, (C) 376.96, or (D) MGA271 CAR T cells. n=5-6 per group.

(E) Mean tumor growth curves for the treatment groups shown in (A). Data shown as mean±SEM.

(F) Kaplan–Meier survival curves corresponding to the study in (A) to (E). Statistical significance was assessed using a log-rank (Mantel–Cox) test. n=5 (UTD, Y111, and MGA271) or 6 (376.96).

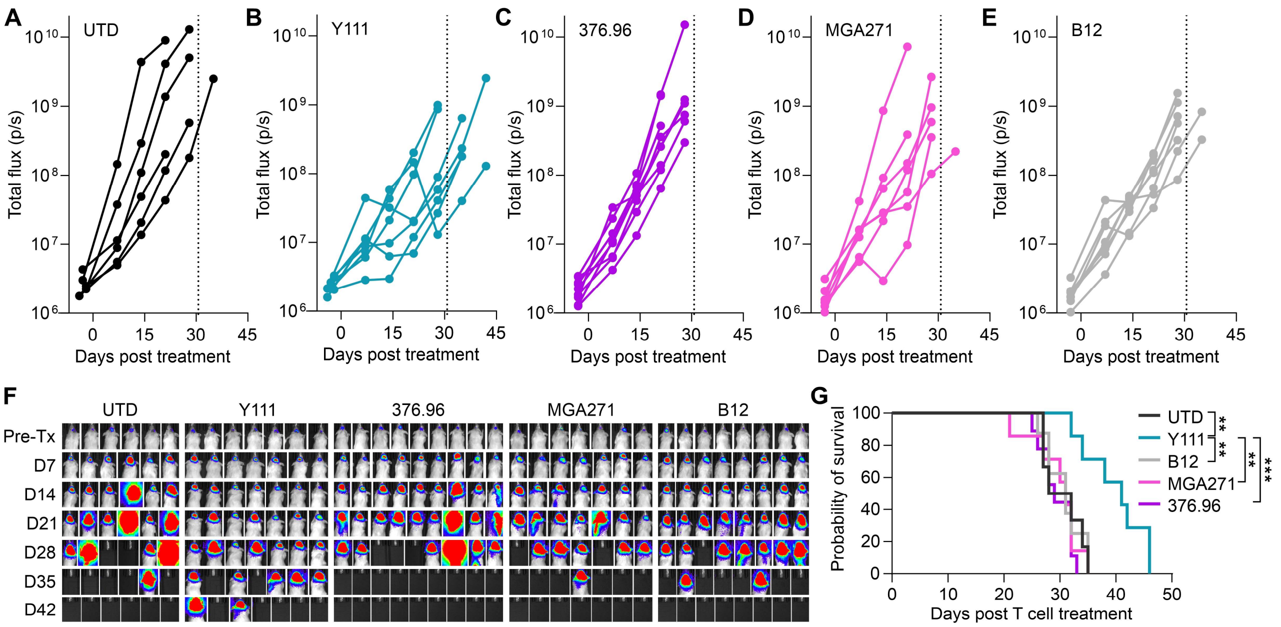


**Supplemental Figure 5. Y111 compares favorably with benchmark B7-H3 CARs in a glioblastoma model.**

(A-E) BLI was used to monitor growth of luciferase labeled GBM01 glioblastoma following stereotactic orthotopic implantation of tumor cells and subsequent i.v. administration of (A) UTD control, (B) Y111, (C) 376.96, (D) MGA271 or (E) B12 CAR T cells. Groups were randomized for treatment based on an equivalent tumor luminescence. Each line represents an independent animal. n=6-9 per group.

(F) Images from the study shown in (A) to (F).

(G) Kaplan–Meier survival curves corresponding to the study in (A) to (F). Statistical significance was assessed using a log-rank (Mantel–Cox) test. n=6 (UTD), 7 (Y111 and MGA271), 8 (B12), or 9 (376.96).

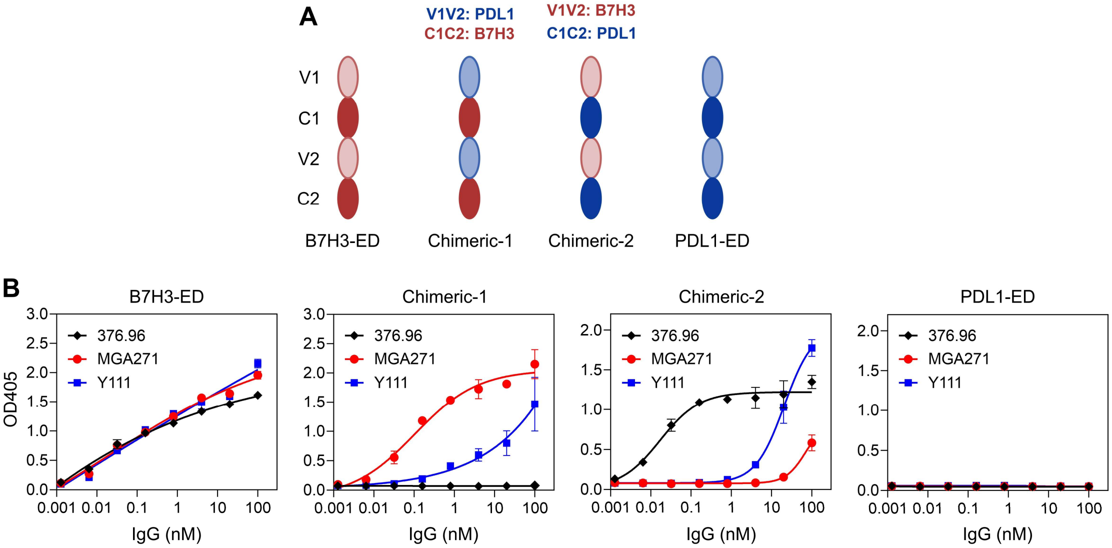


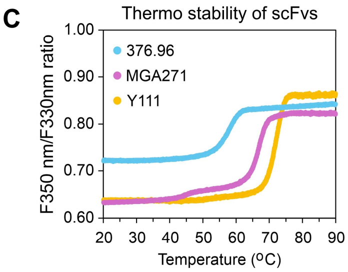


D

Observed composition of B7H3-ED/Ab IgG complex at different input ratios

| mAb | B7H3-ED : full IgG input ratio 1:1 | 1:2 | 1:4 | Ab epitope location |
|---|---|---|---|---|
| 376.96 | 1:2 (73%) | 1:2 (55%) | 2:2 (64%) | IgV domain |
| MGA271 | 2:2 (100%) | 2:2 (100%) | 2:2 (100%) | IgC domain |
| Y111 | 1:1 (100%) | 1:1 (100%) | 1:1 (68%), 1:2 (12%) | IgC and IgV domain |

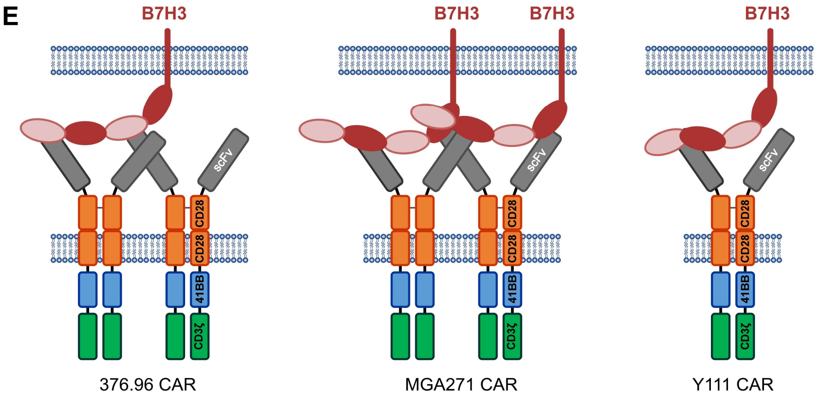


**Supplemental Figure 6. Y111 displays unique binding kinetics.**
(A) Schematic overview of B7-H3 and PDL1 extracellular domain (ED) fusion proteins used in domain-swap experiments to map the binding domain of the 376.96, MGA271 and Y111 antibodies.
(B) ELISA was used to evaluate binding of the 376.96, MGA271 and Y111 full IgG antibodies to the purified fusion proteins depicted in (A).
(C) Thermal stability of the 376.96, MGA271 and Y111 scFvs.
(D) SEC-MALS was used to determine B7-H3-ED/full IgG complex formation following mixture at different input ratios. Note that the bivalent Y111 antibody strongly prefers 1:1 binding even in the presence of an excess of Y111, consistent with the Biacore data of supplemental figure 3A.
(E) Schematic overview showing how the bivalent Y111 CAR may preferentially bind the 4Ig form of B7-H3 - soluble or membrane bound as shown - in a 1:1 complex.
